# Synaptic basis of a sub-second representation of time

**DOI:** 10.1101/2022.02.16.480693

**Authors:** A. Barri, M. T. Wiechert, M. Jazayeri, D. A. DiGregorio

## Abstract

Temporal sequences of neural activity are essential for driving well-timed behaviors, but the underlying cellular and circuit mechanisms remain elusive. We leveraged the well-defined architecture of the cerebellum, a brain region known to support temporally precise actions, to explore theoretically whether the experimentally observed diversity of short-term synaptic plasticity (STP) at the input layer could generate neural dynamics sufficient for subsecond temporal learning. Simulated synaptic input generated a diverse set of transient, firing patterns in granule cells (GCs) that provided a temporal basis set for learning precisely timed pauses of Purkinje cell activity associated with delayed eyelid conditioning and Bayesian interval estimation. The learning performance across time intervals was influenced by the temporal bandwidth of the GC basis, which was determined by the input layer synaptic properties. The ubiquity of STP throughout the brain positions it as a general, tunable cellular mechanism for sculpting neural dynamics and fine-tuning behavior.

## Introduction

The neuronal representation of time on the sub-second timescale is a fundamental requisite for the perception of time-varying sensory stimuli, generation of complex motor plans, and cognitive anticipation of action^1–4^. But how neural circuits acquire specific temporal contingencies to drive precisely timed behaviors remains elusive. A progressive increase in firing rate (“ramping”) towards a threshold can represent different elapsed times by altering the slope of the ramping behavior. Elapsed time can also be encoded by a population of neurons that fire in a particular sequence (“time cells”)^5–8^. Sequential synaptic connections between neurons (synfire chains) can explain the neural sequences representing bird song^9^ and contribute to time delays necessary to cancel self-generated sensory stimuli in the electrosensory lobe of mormyrid fish^10^. Temporal dynamics of neural population activity can also be reproduced by training recurrent neural network models^11–13^. Nevertheless, the search for a candidate neural mechanism for generating a temporal reference (biological timer) for neural dynamics is an ongoing challenge.

STP, the rapid change in synaptic strength occurring over 10’s of milliseconds to seconds, transforms patterned activity in the time domain^14^. Depression and facilitation of synaptic strength can act as low-and high-pass filters, respectively^15^, and synaptic depression can mediate gain modulation^16,17^. Within recurrent neural networks, the long timescales of cortical synaptic facilitation provide the substrate for working memory^18^. Finally, low-gain recurrent networks that are supplemented by STP can also generate neural representations of time^19^. However, experimental evidence of STP-dependent circuit computations is rare, and is largely associated with sensory adaptation^20^.

The cerebellar cortex (CC) is a prototypical microcircuit known to be important for generating temporally precise motor^21^ and cognitive behaviors^22–25^ on the sub-second timescale. Its largely feed-forward circuitry has been proposed to learn the temporal contingencies required for prediction from neural sequences across the population of granule cells (GCs) within the input layer^26^ that excite the sole output of the CC, Purkinje cells (PCs). The synapses between mossy fibers (MFs) and GCs, which convey contextual information to GCs, are highly variable in their synaptic strength and STP time course^27^. Therefore, we hypothesized that STP of MF-GC synapses could be used as internal timers for a population clock within the cerebellar cortex to generate neural dynamics necessary for temporal learning.

To elaborate this hypothesis, we modeled the CC as a two-layer perceptron network that includes realistic MF-GC connectivity and STP dynamics. The model reproduces learned PC activity associated with a well-known temporal learning task: delayed eyelid conditioning^28^. The timescales of STP determined the temporal characteristics of the GC population activity, which in turn defined the temporal window of PC temporal learning. The width of PC pauses scaled proportionally with the learned time intervals, similarly to experimentally observed scalar variability of the eyelid conditioning behavior^29^. Additionally, we found that STP driven GC activity was well suited to implement a Bayesian estimator of time intervals^30^. We propose that within feed-forward circuits, dynamic synapses serve as tunable clocks that determine the bandwidth of neural circuit dynamics and enable learning temporally precise behaviors.

## Results

### Cerebellar Cortex model with STP

The CC can be modeled as a two-layer perceptron that performs pattern separation of static input patterns^31–34^. When temporal processing is considered, CC models are generally augmented by an additional mechanism that generates temporally varying activity patterns in the CC’s GC layer^10,26,35,36^. To test whether heterogeneous MF-GC STP is sufficient to support temporal learning, we implemented STP of the MF-GC synapse in a CC model, hereafter named CCM_STP_, that deliberately omits all other potential sources of temporal dynamics. In particular, the first set of simulations did not consider recurrent connectivity (**Fig 1b**). STP was simulated using a parallel vesicle pool model of the MF-GC synapse, similar to ref^37^. It comprises two readily-releasable and depletable vesicle pools, synaptic facilitation, and postsynaptic desensitization. To reproduce the observed functional synaptic diversity, we set vesicle fusion probabilities (*p_v_*), synaptic pool sizes (*N*), and synaptic facilitation to match the relative strengths, paired-pulse ratios, and transient behaviors across five different types of synapses that were previously characterized^27^ (**Fig 1a_2_-a_6_**). Importantly, the longest timescale in CCM_STP_ is associated with a 2 s vesicle refilling time constant of the slow vesicle pool (*τ_ref_* = 2s, **Fig. 1a_1_**). To capture depression over long timescales^37,38^, we introduced a phenomenological parameter (*p_ref_* = 0.6) that effectively mimics a simplified form of activity-dependent recovery from depression (see Methods).

**Figure 1.**
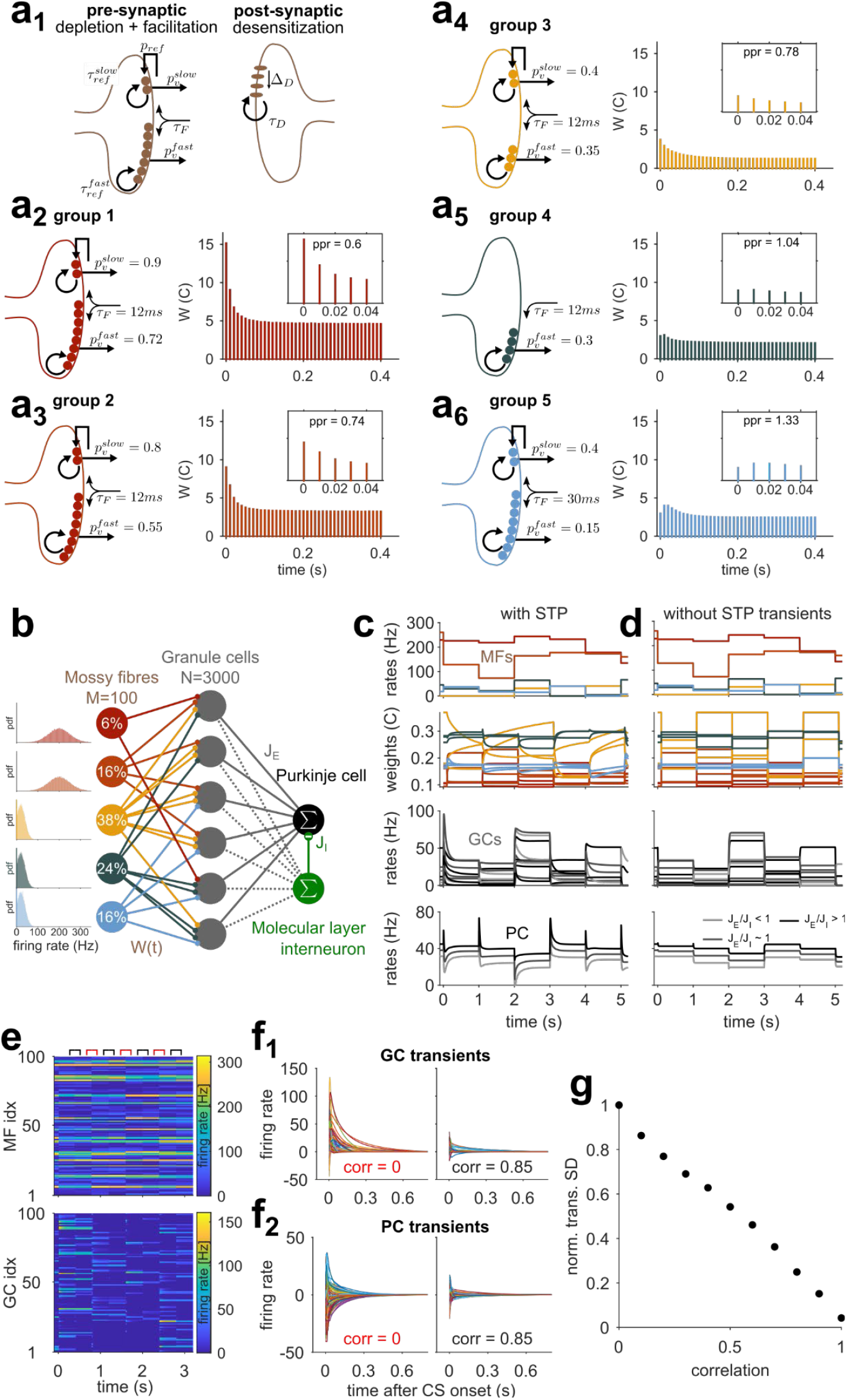
Cerebellar cortex model with dynamic input layer synapses (CCM_STP_) **(a_1_)** Synaptic model scheme. **(a_2-6_)** Properties of five synapse types. Left: Schemes show differences in presynaptic parameters; the postsynaptic side is identical for all groups. Right: average synaptic weights in response to repetitive 100 Hz stimulation as in^38^. Insets: First five responses with paired-pulse ratio (PPR) roughly mimic results from Ref.^27^. Color code for synapse groups is the same as in Ref.^27^. **(b)** Scheme of CCM_STP_. MFs are classified according to the groups in **a**. Percentages indicate relative frequency of MF groups. Insets: firing rate distributions for different MF groups. **(c)** Simulation of CCM_STP_ with randomly drawn J_E_ weights. First panel: 5 sample MFs. Every second, MF activity is re-drawn from distributions in **b**. Second panel: Normalized weights of 10 MF-GC synapses. Third panel: activity of 10 sample GCs. Last panel: PC activity with different shades of gray indicating different E/I ratios onto the PC. **(d)** Same as **c** but without STP transient dynamics. Low amplitude GC and PC firing rate transients result from 10 ms GC integration time constant. **(e)** Example simulation in which correlated (black symbols) and uncorrelated (red symbols) MF patterns were presented to the network in alternation. The correlation coefficient for sequential patterns was ≈ 0.85. Firing rates are color-coded. **(f_1_)** Steady-state subtracted GC responses from simulation in **e** for uncorrelated (left) and correlated MF pattern switches (right). **(f_2_)** Same as **f_1_** but for PC. **(g)** Normalized standard deviation of PC transient amplitudes for switches between MF patterns of differing levels of correlation.

The CCM_STP_ consisted of firing rate units representing MFs, GCs, a single PC, and a single molecular layer interneuron (MLI), i.e. each neuron’s activity was represented by a single continuous value corresponding to an instantaneous firing rate. Each GC received 4 MF synapses, randomly selected from the different synapse types according to their experimentally characterized frequency of occurrence^27^. Importantly, we associated different synapse types with different MF firing rates (**Fig. 1b**, left panels, see Methods). Concretely, we associated synapses with high *p_v_* with MF inputs with comparatively high average firing rates (primary sensory groups 1, 2) and low *p_v_* synapses with MF inputs with comparatively low average firing rates (secondary/processed sensory groups 3, 4, 5), according to experimental observations^39,40^. We will reconsider this relationship below.

To examine CCM_STP_ network dynamics, input MF activity patterns were sampled every second from respective firing rate distributions shown in **Fig. 1b**. Each change in MF patterns evoked transient changes in MF-GC synaptic weights, which in turn generated transient GC firing rate responses that decayed at different rates to a steady-state (**Fig. 1c**). Similar to experimentally recorded PC responses to sensory stimuli *in vivo*^41^, switches between different MF patterns also generated heterogeneous transient changes in the PC firing rate, whose directions and magnitudes were controlled by the ratio of the average excitatory to inhibitory weight (**Fig. 1c, bottom**). In contrast, when MF-GC STP was removed, the transient GC and PC responses disappeared (**Fig. 1d**). The amplitude of the firing rate transients increased as the difference from one MF pattern to the next increased, similar to previous theoretical work^16^. Sequential delivery of uncorrelated MF firing patterns in CCM_STP_ (**Fig. 1e**) generated GC and PC transients with broadly distributed amplitudes (**Fig. 1f_1,2_**), which were progressively reduced as the relative change in MF rate decreased (**Fig. 1g**). Thus, dynamic MF-GC synapses allow both GCs and PCs to represent the relative changes in sensory stimuli.

### Simulating PC pauses during eyelid conditioning

We next explored whether MF-GC STP diversity permits learning of precisely timed PC pauses associated with the prototypical example of a CC-dependent behavior: delayed eyelid conditioning. In this behavior, animals learn to use a conditioned stimulus (CS) to precisely time a blink in anticipation of an aversive unconditioned stimulus (US). The eyelid closure is driven by a preceding decrease in PC firing rates^28,42^ (**Fig. 2a**). As the CS is typically constant until the time of the US and a precisely timed eyelid response can be learned even if the CS is replaced by direct and constant MF stimulation^43,44^, we modeled CS delivery in the CCM_STP_ by an instantaneous switch to a novel MF input pattern that persists over the duration of the CS (**Fig. 2a**). Most GC activity transients exhibited a characteristic rapid increase or decrease in firing rate, followed by an exponential-like decay in firing rate (**Fig. 2c**). In contrast to other models of eyelid conditioning^26^, the activity of most GCs in the CCM_STP_ peaked once during the transient period, occurring shortly (< 50ms) after the CS onset (**Fig. 2c**). However, the distribution of GC firing rate decay times across the population was highly variable with a fraction of GCs showing decay times to 10% of the transient peak as long as 700 ms (**Fig. 2c and 2d**).

**Figure 2.**
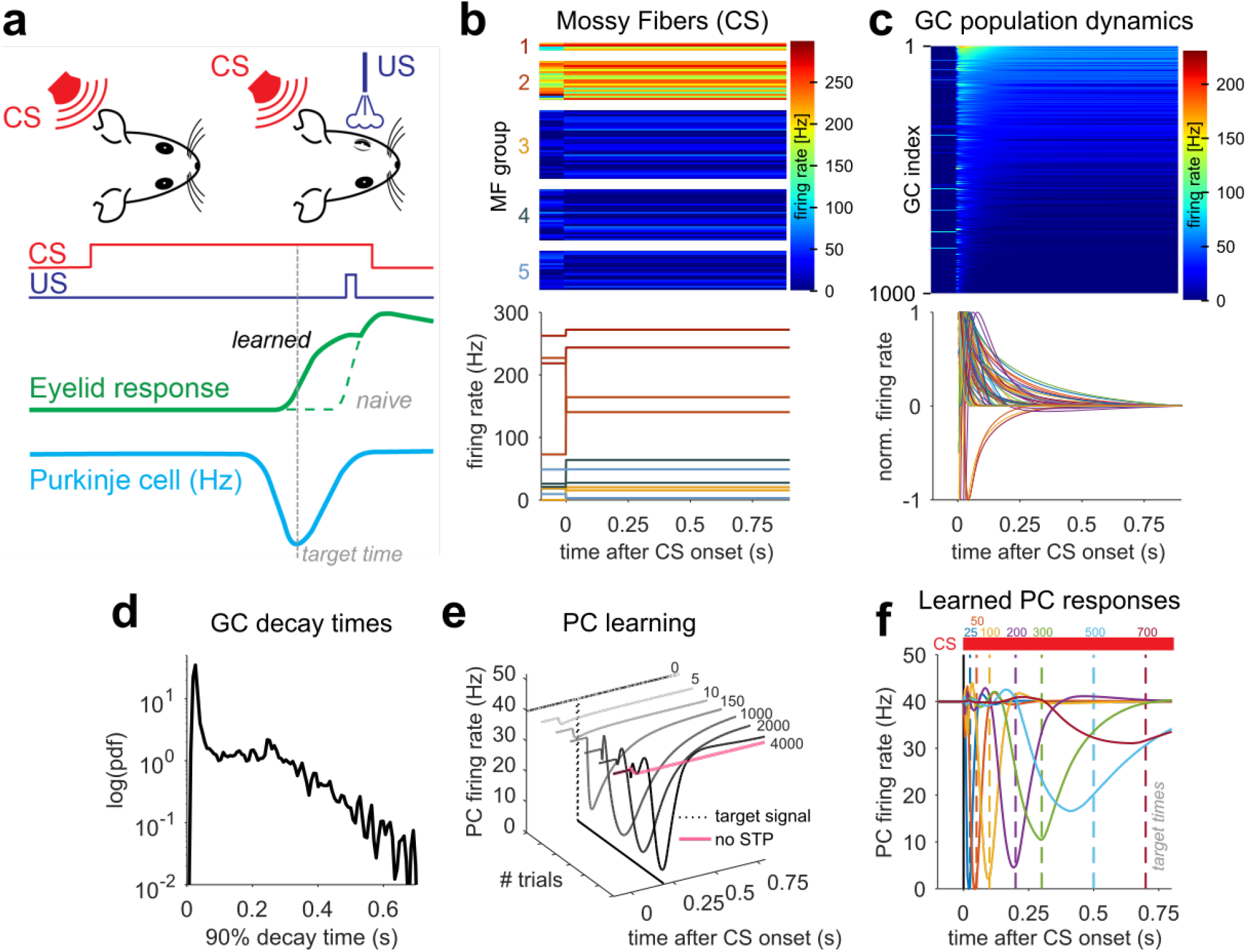
Simulating PC pauses during eyelid conditioning **(a)** Scheme of eyelid conditioning. CS: conditioned stimulus (red). US: unconditioned stimulus (violet). After experiencing CS and US delivered at a fixed temporal contingency over many trials, the animal learns to close its eyelid before the US is delivered (green). A trough in PC activity (blue) precedes the eyelid closure (target time, gray dashed line). **(b)** Representation of the CS by an instantaneous change in MF firing rate. Top: 100 MF sorted according to synaptic types. MF firing rates are color-coded and drawn according to the distributions shown in **Fig. 1b**. Bottom: two sample MF rates per synaptic group. Colors as in **Fig.1**. **(c)** Model GC responses to the CS. Top: 1000 GCs sorted according to average firing rate after CS onset. Firing rates are color-coded. Bottom: steady-state subtracted and individually normalized GC transient responses. **(d)** Pdf of the distribution of GC activity decay times to 10% of the transient peak. **(e)** Example of delayed eyelid conditioning over the course of 4000 learning steps for a 200 ms delay. Dashed line represents the target time used in the supervised learning procedure. Without STP-induced GC transients, no PC pause could be learned (pink line). (**f)** Conditioned eyelid responses for different target times. Different colors indicate PC responses after 4000 learning trials for each target time (dashed lines).

To test whether the GC population dynamics could act as a basis set for learning the precisely timed PC firing rate pauses known to drive the eyelid response, we subjected the GC-PC synaptic weights to a gradient-descent based supervised learning rule. The rule’s target signal consisted of a delta peak (zero firing rate at a specific time bin) at the designated time of the PC firing rate pause (**Fig. 2e**, dotted line). In the course of learning, there was a progressive acquisition of a pause in the PC firing rate (**Fig. 2e**). However, without MF-GC STP, the PC pause was not acquired (**Fig. 2e, pink**). We tested learning of different delay intervals ranging from 25 ms to 700 ms and found that PC pauses could be generated for all delays. The PC pause amplitude and temporal precision (time and width) decreased with increasing CS-US delays (**Fig. 2f**), reminiscent of the shape of PC simple-spike pauses recorded during eyelid conditioning^28^.

Why might the learned PC pause amplitude and temporal precision be reduced for longer CS-US delays? The parameters associated with the learning algorithm (e.g. the number of iterations) are identical for each CS-US delay. The state of the GC population activity, in contrast, changes throughout the CS. Once all GC activity dynamics reach steady-state firing rates, temporal discrimination by PCs is no longer possible, and interval learning becomes impossible. In other words, for temporal learning to be effective, changes in GC firing rates must be prominent over the relevant timescale. Indeed, in delayed eyelid conditioning simulations where slow or fast GCs were removed, the efficiency of generating PC pauses for short and long intervals was reduced (**Fig. S2**). CCM_STP_ simulations thus demonstrate that a GC temporal basis generated by MF-GC STP is sufficient to reproduce the CC computation underlying delayed eyelid conditioning and suggests that the timescale of GC dynamics influences the timescale of behavioral learning.

### Analysis of the synaptic mechanism underlying GC transient responses using a reduced model

PC temporal learning requires GC transient responses, which in our model can only arise from STP at the MF-GC synapse^27,38^. How are synaptic and GC dynamics determined by quantal and firing rate parameters? The complexity of the full CCM_STP_ with many interacting parameters makes it difficult to assess the effect of each synaptic parameter. To overcome this challenge, we developed a reduced MF-GC synapse model which was analytically solvable for an instantaneous and persistent switch of MF rates. This allowed us to identify the key computational building blocks of CCM_STP_ and understand how they control the overall behavior of the model. Specifically, we omitted short-term facilitation and postsynaptic desensitization and reduced the synaptic model to a single population of high *p_v_* synapses (“drivers”^27^) and a single population of low *p_v_* synapses (“supporters”^27^), each with a fast and a slow refilling ready-releasable pool (**Fig. 3b**), thus obtaining a model where STP results from vesicle depletion only. Each GC received exactly two driver and two supporter MF inputs with random and pairwise distinct identities (**Fig. 3a**).

**Figure 3.**
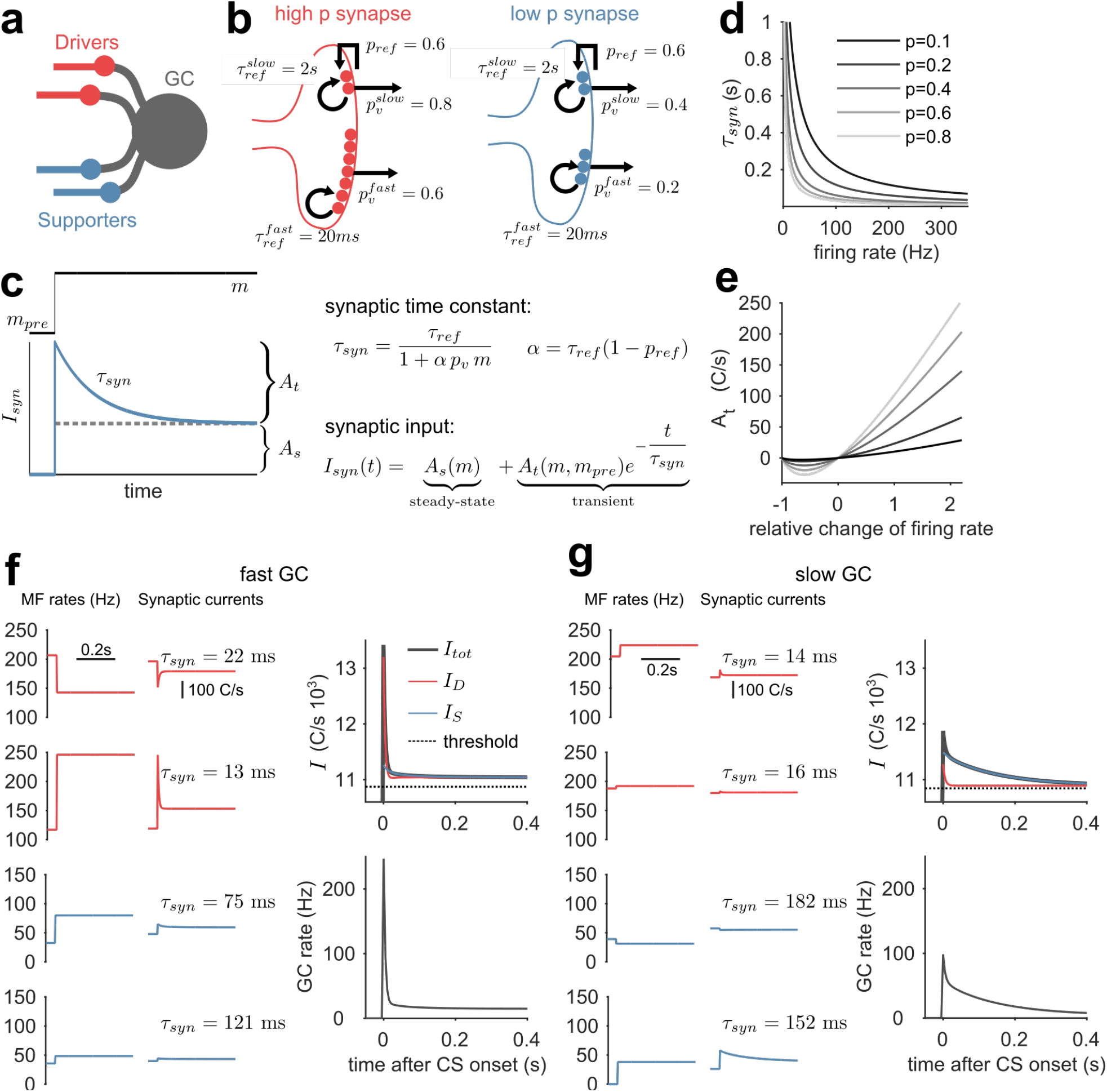
MF-GC synaptic time constants and their relative weight determine the time course of GC responses **(a)** Scheme of GC inputs with the simplified synaptic model. Each GC receives exactly two distinct driver (red) and supporter MFs (blue). **(b)** Schemes of the reduced synaptic model of high p_v_ (red) and low p_v_ synapses (blue). **(c)** Left: scheme of single pool response with the time constant τ_syn_ (blue line) to a firing rate switch during CS presentation (black solid line). The dashed black line separates the transient (A_t_) from the steady-state amplitude (A_s_). Right: equations determining the synaptic time constant and synaptic input. **(d)** Slow vesicle pool time constant (τ_syn_) versus presynaptic MF firing rate. Different shades of gray indicate different release probabilities. **(e)** Driver synapse transient amplitude (A_t_) versus relative firing rate change for a baseline firing rate of 80 Hz ((m – m_pre_)/m_pre_, m_pre_ = 80 Hz). A negative A_t_ corresponds to a transient decrease in firing rate. Same color code as in **d**. **(f)** Sample fast GC. Left: driver and supporter MF firing rates drawn from thresholded normal distributions (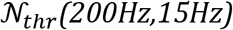 and 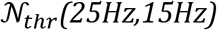, respectively) and the corresponding synaptic responses. For clarity, only the τ_syn_ of the respective slow pool is indicated. Upper right panel: GC threshold (dashed line), total synaptic input (black line), total driver input (red line), and total supporter input (blue line). The transient response is dominated by the driver input (red). Lower right panel: resulting GC firing rate response. **(g)** Like **f** but for a sample slow GC. The transient response is dominated by the supporter input (blue).

In this reduced model, an instantaneous and persistent switch of MF rate change generates an average postsynaptic current (*I_syn_* (*t*)) for each vesicle pool that is remarkably simple. It features a sharp transient change, followed by a mono-exponential decay to a steady-state synaptic current amplitude, *A_s_*, (**Fig 3c**) and can be generally expressed as:

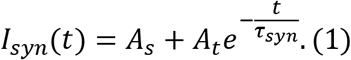

Here, *A_s_* is time-invariant, and 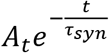 is a time-dependent component with synaptic relaxation time constant *τ_syn_* (**Fig 3c**) and amplitude *A_t_*. This transient component determines the synapse’s ability to encode the passage of time

The solution of the synaptic dynamics model reveals the crucial dependence of *τ_syn_* and *A_t_* on the presynaptic and firing rate parameters (see Methods):

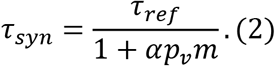

Here, *α* = *τ_ref_*(1 – *p_ref_*), *m* is the MF firing rate persisting during the CS, and the synaptic parameters *p_v_*, *τ_ref_*, and *p_ref_* are defined as above. Equation (2) shows that *τ_syn_* is inversely related to the MF rate during the CS and the release probability, *p_v_* (**Fig. 3d**). Intuitively, this is because higher *p_v_* and/or *m* lead to a higher rate of synaptic vesicle fusion, and hence depletion, driving the synaptic response amplitude to steady-state faster. Conversely, slow time constants arise from low *p_v_* and/or low *m* with the maximum *τ_syn_* being equal to the vesicle recovery time *τ_ref_*.

The transient amplitude *A_t_* is given by:

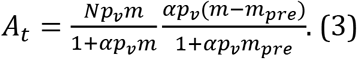

Here, *N* is the number of release sites. Importantly, and in contrast to *τ_syn_*, *A_t_* also depends on the presynaptic MF firing rate before the CS, *m_pre_*, and on the difference between the MF firing rates before and during the CS. In particular, for both rates sufficiently high, *A_t_* becomes a linear function of the normalized difference between *m* and *m_pre_*, i.e. *A_t_* ∝ (*m* – *m_pre_*)/*m_pre_* (**Fig. 3e**). *A_t_* is sensitive to the relative and not the absolute change in presynaptic rate, as observed previously^16^.

The transient GC activity results from the sum of eight synaptic transient current components, (i.e. four inputs, each with two pools). To illustrate the interplay between the *A_t_* and *τ_syn_*, we compared the behavior of each synaptic input for a selected fast and slow GC (**Figs. 3f** and **3g)**. Generally, synaptic inputs from supporters display longer transient currents than synaptic inputs from drivers (**Figs. 3f and 3g**, middle panels) due to their lower firing rates (**Figs. 3f and 3g**, left panels) and low *p_v_* (**Fig. 3b**). *A_t_* is largely determined by the relative change in the respective presynaptic MF rates, (*m* – *m_pre_*) */m_pre_* (Figs **3f** **and 3g, left panels**). Thus, as driver synapses are stronger than supporter synapses, “fast” GCs are generated when driver inputs exhibit large relative changes in firing rates (**Fig.3f**). “Slow” GCs are generated from synapses with a small relative change in driver firing rates, but large relative supporter rate changes paired with low supporter rates during the CS (**Fig.3g**). Taken together, in the reduced model *τ_syn_* and *A_t_* determine the effective timescales of the GC firing rate responses and are explicitly influenced by quantal parameters, synaptic time constants, and the diversity of MF firing rates.

### The explicit influence of synaptic parameters on temporal learning

Our simulations suggest that delayed eyelid conditioning across multiple delays necessitates GC population dynamics spanning multiple timescales (**Fig. 2**, **Fig. S2**). Since individual GC firing rate dynamics depend on the *A_t_* and *τ_syn_* of their synaptic inputs (**Fig. 3**), this implies that, firstly, the spectrum of *τ_syn_* available to the network should cover the relevant timescales, and secondly, the *A_t_*, which can be understood as the relative weights of synaptic transient components, should be of comparable magnitude across *τ_syn_*. To illustrate these points, we used the reduced CCM_STP_ to simulate eyelid response learning with different firing rate properties and examined the relationship between *τ_syn_*, *A_t_*, the GC temporal basis, and learning outcome. Importantly, since *A_t_* and *τ_syn_* are not independent, the quantity of interest is their joint distribution. We initially set up a reference simulation by choosing MF firing rate distributions such that the diversity of GC transient responses and the temporal learning performance (**Fig. 4a**) were comparable to the CCM_STP_ with native synapses (**Fig. 2f**). For this case, the joint distribution shows that *A_t_* decreased with increasing *τ_syn_*. Note that *A_t_* is maximal when the MF firing rates increased from zero *m_pre_* to a finite *m* upon CS onset, maximizing *m-m_pre_* (eqn. 3, see also **Fig. S3b,c**). We quantified learning accuracy by calculating an error based on 1) the PC response amplitude, 2) its full width at half maximum and 3) the temporal deviation of its minimum from the target delay (**Fig. 4a**, fifth panel, **Fig. S2a**, see Methods). Importantly, the degradation in temporal precision of the learned PC pauses for longer CS-US delays was concomitant with the reduction of the *A_t_* associated with longer *τ_syn_* (**Fig. 4a**). This suggests that inspection of the joint distribution of *τ_syn_* and *A_t_* can provide insight into the temporal learning performance of the network.

**Figure 4.**
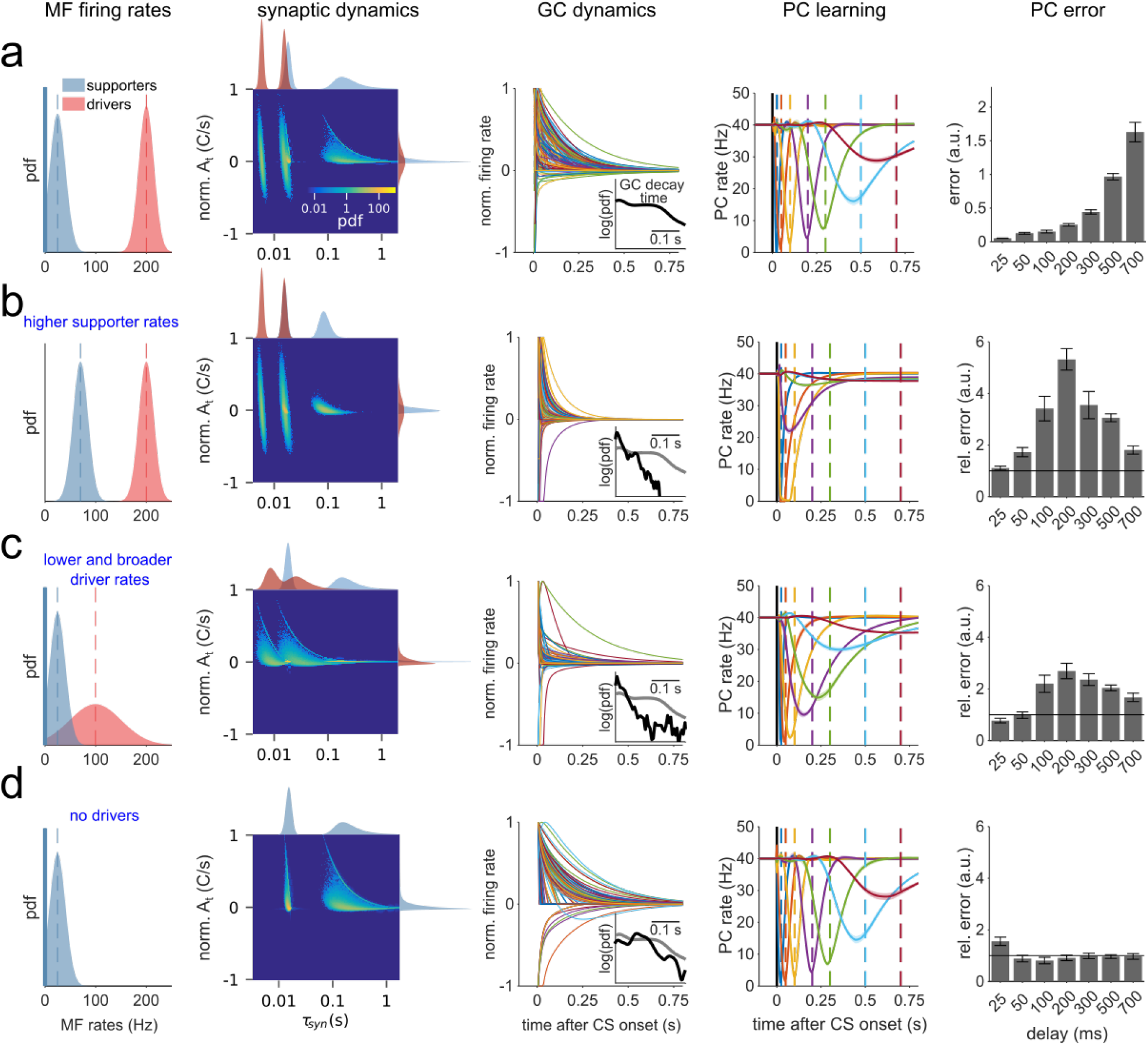
Learning performance depends on MF rate distributions **(a)** First panel: Driver and supporter MF firing rate pdfs (μ_D_ = 200Hz, μ_s_ = 25 Hz, σ_D_ = σ_s_ = 15Hz). Second panel: Resulting joint A_t_ and τ_syn_ distribution, featuring four partially overlapping clusters, corresponding to the slow and fast pools for driver and supporter synapses, and marginal distributions. The color code of the joint distribution scales logarithmically. Colors of marginal distributions indicate driver (red) and supporter (blue) components. Third panel: Normalized GC transient responses to CS. Inset: pdf of the distribution of decay times to 10% of the transient peak. Fourth panel: PC eyelid response learning. Color code delay intervals as in (**Fig. 2e**). Fifth panel: Error for each delay, calculated based on PC response amplitude, full width at half maximum and temporal deviation (**Fig. S2a**). **(b)** Same as **a**, but with μ_s_ = 70Hz. Inset: black line is the pdf for simulation with μ_s_ = 70Hz and gray line is the pdf from **a** for comparison. Fifth panel: errors relative to the ones in **a**. **(c)** Same as **a**, but with μ_D_ = 100Hz and σ_D_ = 50Hz. **(d)** Same as **a**, but without driver inputs.

When changing only the mean firing rate of *supporter* MFs (*μ_s_*) from 25 Hz to 70 Hz, the synaptic time constants were shortened due to the inverse relationship between *τ_syn_* and the mossy-fiber firing rate *m* (**Fig. 4b**, second panel). Consequently, and expectedly, the distribution of GC firing rate decay times was shifted to shorter values, and learning performance was degraded for all CS-US intervals, except the 25 ms delay (**Fig. 4b**). Lowering the mean firing rate of *driver* MFs (*μ_D_*) from 200 Hz to 100 Hz and increasing the standard deviation (*σ_D_*) from 15 Hz to 50 Hz, led to an overall increase of the time constants contributed by driver synapses, as well as an increase in their relative weight (*A_t_*; **Fig. 4c**, second panel, marginals). As a result the joint probability distribution shows a shift towards faster weighted time constants. It also follows that GC transients are accelerated, and learning precision is decreased. Removing synaptic currents originating from driver synapses only disrupted learning PC pauses for the shortest CS-US interval (**Fig. 4d**). Reduced model simulations with systematic parameter scans across a wide range of mossy fiber firing rate distributions for both synapse types suggested that good synaptic regimes for temporal learning are achieved when driver synaptic weights are comparable or smaller than those of the slow supporting synapses (**Fig. S4**).

All the results taken together suggest that optimal learning occurs when the spectrum of *τ_syn_* available to the network covers behaviorally relevant timescales with balanced relative weights (*A_t_*). Synaptic and GC activity timescales can therefore be tuned by simultaneously modulating *p_v_* and the absolute scale of *m* to provide the necessary distribution of *τ_syn_*, whereas the relative change of MF firing can be used to tune the weights (*A_t_*) of *τ_syn_*.

### Firing rate and synaptic parameters that improve temporal learning performance

Thus far, we used the reduced model to explore how MF firing rates and synaptic properties influenced the timescales of GC activity and the temporal precision of learned PC pauses. The model, however, was constrained by 1) the use of only two synapse types, 2) fixed release probabilities (*p_v_*), 3) MF firing rates that were consistently higher for high *p_v_* synapses than their low *p_v_* counterparts, and 4) an equal number of driver and supporter synapses. We next considered how the relaxation of these assumptions and specific parameter combinations could influence the precision of learned PC pauses. In particular, we simulated reduced models where, in addition to MF firing rates, *p_v_* was sampled from continuous distributions.

Equation 2 suggests that a positive correlation between *p_v_* and m should broaden the distribution of *τ_syn_* and broaden the time window of learning. Specifically, we expect learning performance to improve when high(low) firing rate MFs are, on average, paired with high(low) *p_v_* synapses. We chose uniformly distributed *p_v_* and MF firing rates and split both of these equally into two contiguous groups (**Fig. 5a**). We performed training simulations in which we paired high *p_v_*(driver) synapses with high firing rates, or we paired low *p_v_* (supporter) synapses and high MF firing rates, and vice versa (**Fig. 5b**). Formally, this is equivalent to adjusting the rank correlation (*c_rk_*) between the *p_v_* category (supporter or driver) and the *m* category (high or low, Fig. 5b). We found better learning performance when *p_v_* and *m* were positively correlated (**Fig. 5c, Fig. S5**), consistent with the experimentally observed high firing rates for the driver-like synapses arising from primary vestibular afferents^27,39^ and low firing rates for the supporter-like secondary vestibular afferents^27,40^.

**Figure 5.**
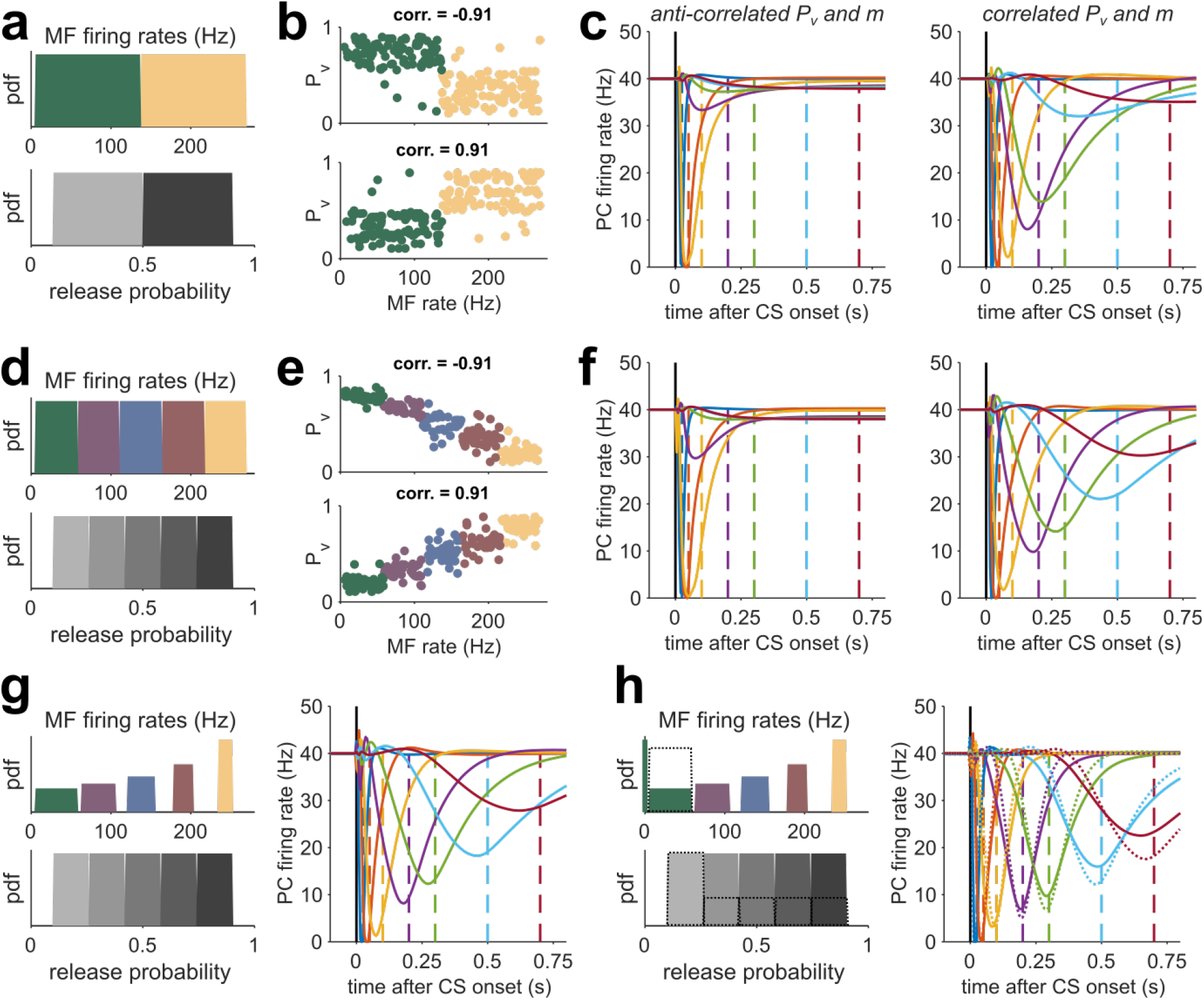
Correlating release probability and MF rates improves learning performance **(a)** Top: distribution of MF firing rates (m) used to drive the network, divided into low (supporter, green) and high (driver, yellow) rates. Bottom: Distribution of synaptic release probabilities (p_v_), divided into low (light gray) and high (dark gray) probabilities. **(b)** Top: p_v_ versus m for 500 sample synapses for a network with strong negative rank-correlation between the m category (supporter or driver) and the p_v_ category (high or low). Bottom: same as top, but for strongly positive correlated m and p_v_. **(c)** Resulting PC eyelid response learning for low (left) and high (right) correlation, with delay intervals color-coded as in (**Fig. 2f**). Each curve is the average of 20 network realizations. **(d-f)** Same as **a**, but for distributions divided into five groups. **(g)** Left: MF rates and release probabilities for 5 groups where the average group firing rate is as in **d** but the firing rate variance progressively decreases with the average rate. Right: resulting PC eyelid response learning for high correlation. **(h)** Same as **g** but with zero rate MFs added to the lowest rate distribution. Dashed lines indicate additionally the case when the count of lowest rates and release probabilities is doubled.

Inspired by the number of synapse types observed experimentally^27^, we augmented the number of synapse groups from 2 to 5 without changing the *p_v_* and firing rate distributions (**Fig. 5d**). We reasoned that the introduction of a larger number of MG-GC synapse types would in principle permit a stronger *linear* correlation between *p_v_* and *m* to occur (**Fig. 5e**), leading to a broader *τ_syn_* spectrum (not shown) and an improved learning of PC pauses. Indeed, for high *c_rk_*, the learning performance of the five group CCM_STP_ was better than that of the two-group CCM_STP_ (compare **Fig. 5c** and **Fig. 5f, Fig. S5**). These simulation results suggest that good temporal learning performance of CCM_STP_ can be achieved not simply by generating variability in parameters, but by structuring, or tuning, the relationship between *p_v_* and *m*.

Equipped with an understanding of how the synaptic and MF rate parameters can generate different synaptic time constants, we set out to further improve the temporal learning for longer CS-US delays by adjusting the variance of the clustered MF rate distributions. To increase the weighting of long *τ_syn_*, we inversely scaled the variance of the MF firing rate distributions with respect to the mean firing rate (**Fig. 5g**), thereby increasing *A_t_* (**Fig. 4c**). As expected, PC pause learning was better than when using equal-width MF groups (**Fig. 5g**, **Fig. S5**). An additional enhancement of learning performance could be achieved by adding a small fraction of zero-rate MFs to the lowest group (**Fig. 5g**, 6% zero MFs, same fraction as in **Fig. 4a**), which provide maximal *A_t_* (see **Fig. 4**). Finally, taking into account the experimental finding that low *p_v_* synapses are more frequent than high *p_v_* synapses^27^, we doubled the fraction of MFs and release probabilities in the lowest group, resulting in the best performance of all versions of CCM_STP_ tested here (**Fig. 5g**). These simulations show that positive correlations between vesicle release probability and presynaptic firing rate broaden the temporal bandwidth of circuit dynamics and improve temporal learning.

### STP permits learning optimal estimates of time intervals

Humans and animals have an unreliable sense of time. When estimating or producing time intervals, behavioral response variability scales linearly with the base interval^45^. Previous work has found that humans seek to optimize their timing behavior and minimize the effect of scalar variability. A canonical example of this optimization is evident in the so-called ready-set-go-task^46^ (**Fig. 6c**). In this task, subjects must estimate and subsequently reproduce different time intervals. When the intervals are drawn from a previously learned probability distribution (i.e., prior), subjects generate biased responses that are in accordance with an optimal Bayesian agent. Bayesian integration predicts that when the prior distribution is uniform, time interval production is biased towards the mean of the prior distribution, and that this bias is larger for longer intervals.

**Figure 6.**
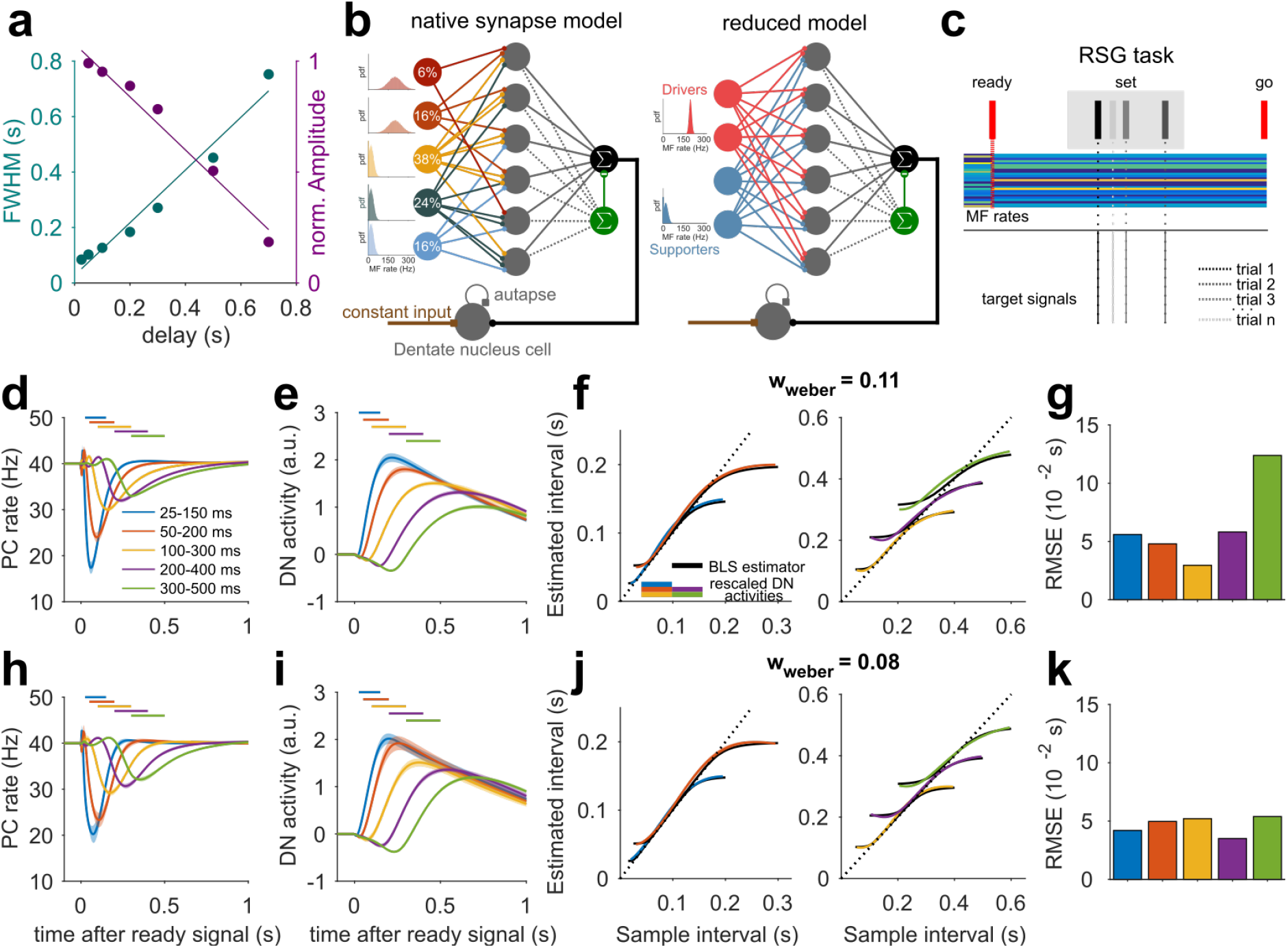
STP generated temporal basis enables the computation of Bayesian time-interval estimates **(a)** Full width at half maximum (cyan) and normalized amplitudes (magenta) of learned PC eyelid responses versus delay interval from the experimentally constrained CCM_STP_ (see **Fig. 2**). Solid lines are linear fits. **(b)** Schemes of native synapse CCM_STP_ (left, see **Fig. 1**) and reduced model (right) with added dentate nucleus cell. **(c)** Scheme of ready-set-go task. The interval between ready and set (sample interval) is drawn from a uniform distribution at every learning iteration (gray box). The ready signal is represented by an instantaneous and persistent switch in MF firing rates (colored lines). The target signal is changed according to the set interval (dotted lines). **(d)** PC responses after 12000 iterations of RSG learning, averaged over 20 network realizations. Shaded area indicates standard deviation. Different colors represent learning of different uniform sample interval distributions. **(e)** Same as **d**, but for DN cell activity. **(f)** Rescaled DN cell activity for different learnt interval distributions (colored) and fitted theoretical Bayesian least squares (BLS) estimator (solid black line), with w_weber_ = 0.11 resulting from fit. **(g)** Squared deviation of rescaled DN activity from the BLS estimator for all tested intervals. **(h-k)** Same as **d-g**, but for the reduced model. The reduced model firing rate parameters were μ_D_ = 200Hz, μ_s_ = 20 Hz, σ_D_ = 10Hz and σ_s_ = 15Hz and resulted in DN activity was consistent with a Bayesian least squares model with w_weber_ = 0.08.

A recent study developed a CC circuit model called TRACE that could emulate the Bayesian computations during the ready-set-go task^30^. In TRACE, learning is mediated by long-term synaptic plasticity of the GC-PC synapses in subsets of GCs that are active at the time of “set.” After learning, PC firing rates undergo prior-dependent pauses that enable the downstream dentate nucleus neurons (DNs) to represent a Bayesian estimate of the time interval. An essential feature of TRACE is that the GCs form a temporal basis set that exhibits temporal scaling. This temporal scaling, which is inherited by PCs, enables DNs to generate estimates that take into account the inherent scalar variability of timing.

In our analysis of eyelid conditioning (**Fig. 2**), we showed that CCM_STP_ generates PC firing rate pauses whose width and amplitude scale with time (**Fig. 6a**). Therefore, we reasoned that CCM_STP_ might also be able to generate biased responses consistent with Bayesian integration. To test this possibility, we analyzed the behavior of CCM_STP_ in the ready-set-go task. We treated the “ready” signal as an instantaneous switch of the MF input rates that persisted over the course of a trial (**Fig. 6c**), similar to the CS in the eyelid simulations. To learn a distribution of ready-set intervals, we generated target signals with intervals that varied from iteration to iteration of our learning algorithm (**Fig. 6c**).

We tested CCM_STP_ with five different uniform distributions of ready-set intervals (25-150 ms, 50-200 ms, 100-300 ms, 200-400 ms, 300-500 ms), resulting in PC pauses that broadened for longer interval distributions, and integrated DN activity that could easily match the Bayesian least-square model^30^ by adjusting a single parameter, the Weber fraction *w_weber_* (see Methods; **Fig. 6d,h**). Overall, the reduced model interval estimates were closer to the Bayesian optimal estimates than for CCM_STP_ with native synaptic parameters, especially for the 200-400 ms and 300-500 ms intervals (**Fig. 6h-k**). These simulations show that a GC basis generated by MF-GC STP is sufficient for driving Bayesian-like learning of time interval estimates on timescales of several hundreds of milliseconds. It should be noted that our GC temporal basis was not explicitly constructed to accommodate scalar properties. Nevertheless, as in the TRACE model, we observed that interval estimates were biased towards the mean and that these biases were larger for longer intervals. These results suggest that a GC basis set generated from the diverse properties of native MF-GC synapses likely exhibits a scalar property necessary for generating optimal timed behaviors.

## Discussion

In order to generate temporally precise behaviors, the brain must establish an internal representation of time. This theoretical study posits that the diversity of synaptic dynamics is a fundamental mechanism for encoding sub-second time in neural circuits. By using eyelid conditioning as a benchmark task for the CCM_STP_, we elucidated the conditions under which the variability in MF-GC synaptic dynamics generates a GC temporal basis that represents elapsed time and that is sufficient for temporal learning on a subsecond scale. According to David Marr’s levels of analysis of information processing systems^47^, our study connects all three levels, from the circuit computation (learning timed PC pauses) to its underlying algorithm (learning with a temporal basis set), and the fundamental biological mechanism (STP).

### STP diversity as a timer for neural dynamics

Cerebellar adaptive filter models posit that GCs act as a heterogeneous bank of filters that decompose MF activity into various time-varying activity patterns - or temporal basis functions - which are selected and summed by a synaptic learning rule at the PC to produce an output firing pattern that generates behaviors that minimize error signals arriving via climbing fibers^35,36^. CCM_STP_ can be viewed straightforwardly as an adaptive filter in which MF-GC synapses act as non-linear elements whose filter properties are determined by the experimentally defined synaptic parameters and modulated by the presynaptic MF firing rates. Because of this filter bank, GCs and PCs in CCM_STP_ respond with exponential-like activity patterns to step-like changes in MF firing rates. Interestingly, hippocampal neural sequences can be described using timing models that use a bank of leaky integrators and produce a family of transient neural activity changes that exhibit variable exponential decays in response to instantaneous changes in input firing rates^48^.

The use of an instantaneous and persistent change in MF activity was motivated by the fact that eyelid conditioning can be achieved if the CS is replaced with a constant MF stimulation^43,44,49^. Recent evidence from pons recordings during reaching, suggests that MF activity can be persistent with little dynamics^50^. For dynamic changes in MF rates STP is likely to generate outputs that are phase-shifted and/or the derivatives of their input^51^. Using heterogeneity of MF-GC STP as a mechanism for adaptive filtering, even time-varying inputs will effectively be diversified within the GC layer and improve the precision of temporal learning.

Synapses within the prefrontal cortex^52^ and at thalamocortical connections^53^ exhibit diverse firing rate inputs^54^ and release probability, and would generate synaptic dynamics that could drive complex neural dynamics. Reminiscent of PC firing rate pauses during eyeblink conditioning, hippocampal time cells are thought to be generated by a linear combination of exponentially decaying input activity patterns from upstream entorhinal cortical neurons^6^. We, therefore, speculate that STP diversity could underlie the generation of temporal context-like cells in the cerebellum (GCs) and other brain regions.

### Timing mechanisms in the cerebellar cortex

In addition to MF-GC STP, the cerebellar cortex is equipped with multiple mechanisms potentially enabling temporal learning^55^. Indeed, time-varying MF inputs could directly provide a substrate for learning of elapsed time, but whether the observed diversity of MF firing is sufficient to mediate temporally precise learning is unknown and merits further exploration. Within the cerebellar cortex, unipolar brush cells are thought to provide delay lines to diversify GC activity patterns^10,56,57^, but these cell types are rare outside the mammalian vestibular cerebellum. The diversity of GC STP^58^ could add to the diversity of the effective GC-layer basis set. Computer simulations have shown that recurrent GC-GoC-cell network dynamics could also generate a rich GC temporal basis set^12,26^. However, experimental support of such circuit dynamics is lacking. When we included recurrent GC-GoC synapses in our simulations, more GCs showed a delayed onset firing (**Fig. S6**), but the synaptic time constant distribution and learning rate remained unaltered. These simulations are consistent with a “common mode” GoC cell population activity^59^. Consistent with the importance of MF-GC STP, delayed eyelid conditioning was selectively altered as a result of the loss of fast EPSCs in AMPAR KO mice^60^. Simulations including realistic NMDA, and spillover dynamics^61^ can further enrich the temporal scales available to the network^62^. MF-GC STP and other timing mechanisms described above are not mutually exclusive. They could act in concert with the diverse intrinsic properties of GCs^63^ and PCs^64^ in order to cover different timescales or increase mechanistic redundancy.

### Predictions of the CCM_STP_

Our theory makes several testable predictions. The transient response amplitude of PCs, which is proportional to the relative change in firing rate, can serve as a detector of rapid changes in MF firing patterns (novelty) and thus amplify pattern discrimination similar to that demonstrated for synapse-dependent delay coding^27^. Consistent with this prediction, single whisker deflections have been shown to generate transient PC activity^41^.

The CCM_STP_ is one of the few network models to directly link quantal synaptic parameters and presynaptic activity dynamics to population activity dynamics and temporal learning. In Figures 3 and 4, we showed that the relative weight and temporal span of synaptic time constants dictate the distribution of GC firing rate decay times and, in turn, the timescales of temporal learning. Analytical solutions for simple synapse models (eqn. 3) provide insight into how synaptic parameters influence STP. For example, high levels of correlation between *p_v_* and *m*, coupled with balanced relative weights of the synaptic time constants, generated a learning performance superior to the native synapses (**Fig. 5d**). Therefore, CCM_STP_ predicts that MFs forming driver synapses (high *p_v_*) would have high baseline and stimulated firing rates, while MFs forming supporter synapses (low *p_v_*) would exhibit low baseline and stimulated firing rates, albeit with large relative changes in firing rates. Indeed, vestibular neurons, which have been shown to exhibit high firing rates^65,66^, produced MF-GC synapses with high release probability^27^. In the C3 zone of the anterior lobe in cats, specific firing rates were associated with different MF types^67^. It is tempting to hypothesize that nature tunes presynaptic activity and synaptic dynamics (perhaps by homeostatic or activity dependent mechanisms) in order to preconfigure the window of temporal associations required for a particular behavior.

### Synaptic implementation of a Bayesian computation

Bayesian theories of behavior provide an attractive framework for understanding how organisms, including humans, optimize time perception and precise actions despite the cumulative uncertainty in sensory stimuli, neural representations, and generation of actions^46,68^. We found that CCM_STP_ was capable of generating biased time estimates consistent with Bayesian computations. In general, the magnitude of biases for a Bayesian agent depends on the magnitude of timing variability (i.e., Weber fraction). In our simulations, model parameters that correspond to native synapses from the vestibular cerebellum produced biases that were optimal for a typical weber fraction of 0.11. However, CCM_STP_ is flexible and can be adjusted to generate optimal biases for a wide range of weber fractions. The exact relationship between model parameters and *w_weber_* is an important question for future research. We note that the timescales of synaptic properties observed empirically in the vestibular cerebellum^27^ are only suitable for generating optimal estimates for relatively short time intervals. Therefore, it remains to be seen whether the synaptic mechanisms that underlie CCM_STP_ could accommodate timing behavior for longer timescales. In this respect, one intriguing hypothesis is that synaptic parameters in different regions of the cerebellum are tuned to generate optimal estimates for different ranges of time intervals, similar to the timing variability observed for cerebellar long-term synaptic plasticity rules^69^.

## Online Methods

### MF-GC synapse model

The synaptic weight between the *j*th MF and the *i*th GC is denoted by *W_ij_*. The firing rate of the *j*th MF is represented by *m_j_*(*t*) and the average current per unit time transmitted by the synapse between GC *i*. and MF *j* is:

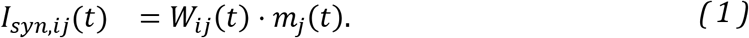

Time-dependent MF-GC synaptic weights were modeled using two ready-releasable vesicle pools^37^, each according to the general form established by Tsodyks and Markram^70^. A similar model was shown to describe accurately STP at the MF-GC synapse^37^. Accordingly, one vesicle pool was comparatively small with high release probability and a low rate of recovery from vesicle depletion (0.5s^−1^) while the other is comparatively large, with low release probability and a high rate of recovery from depletion (20ms^−1^)^37^. We refer to these pools as ’slow’ and the ’fast’, respectively. In the Hallermann model^37^, the slow pool is refilled by vesicles from the fast pool. Here, for the sake of mathematical tractability, we modeled the pools as being refilled independently (see scheme in **Fig. 1**).

To model vesicle depletion we use the variables *χ^slow^* and *χ^fast^*, denoting the fractions of neurotransmitter available at the slow and fast vesicle pool, respectively. The state of the pools between GC *i* and MF *j* at time *t* are then described by:

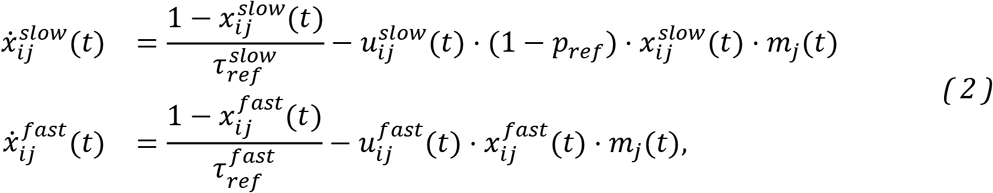

where, 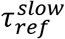 and 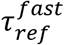 are the time-constants of recovery from vesicle depletion; these are identical across all synapses. The variables 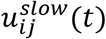 and 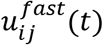 denote the pools’ respective release probabilities at time *t*. Experimental data show that, in response to trains of action potentials, MF-GC synapses approach synaptic steady state transmission with a long time-constant^37,38^, a feature that can be captured with a serial pool model^37^ (see scheme in **Fig. S7**). In order to capture this behavior with a parallel pool model, we added the phenomenological parameter *p_ref_* to the slow pool’s dynamical equation. In mechanistic terms, *p_ref_* can be thought of the probability of immediately refilling a synaptic docking site after the release of a vesicle. This mechanism effectively mimics a simplified form of activity-dependent recovery from depression. The release probabilities 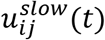 and 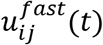 are modulated by synaptic facilitation according to:

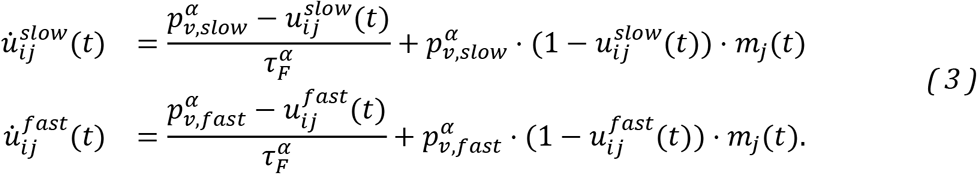

Here, 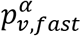 and 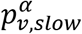 denote the release probabilities for the fast and slow pools, respectively, and 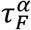 is the facilitation time constant. The index *α* denotes different synaptic groups and runs from 1 to 5. With the above, the average number of vesicles released at any time t can be written as:

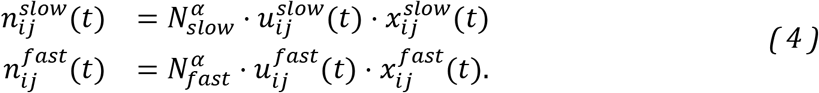

Post-synaptic receptor desensitization induces an additional component of depression of phasic MF-GC synaptic transmission. As both pools share the same post-synaptic side, we model desensitization via the modulation of a single variable *q_ij_* (*t*) which represents the synaptic quantal size and which is influenced by the total number of vesicles released from both pools:

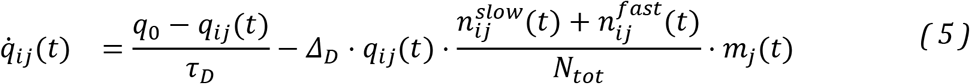

where 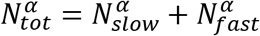 is the time constant of recovery from desensitization, *q*_0_ is the quantal size in absence of ongoing stimulation and *Δ_D_* is a proportionality factor that determines the fractional reduction of *q_ij_*(*t*). As explained below, we set *q*_0_=1, i.e. *q_ij_*(*t*) is normalized. Both *τ_D_* and *Δ_D_* are identical across all synapses. Finally, the total synaptic weight is equal to the sum of the contributions from both vesicle pools:

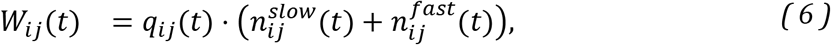

### Synaptic parameters for generating diverse synaptic strength and dynamics

We set the synaptic parameters of our model such as to reproduce the average behavior of the 5 MF-GC synapse groups which were determined in ref^27^ based on unitary response current amplitudes, pair pulse ratios and response coefficients of variation.

The vesicle pool refilling time constants 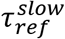 and 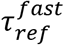 were set to the values measured at the MF-GC synapse in ref.^37^ and were identical for all synapse groups. The time constant of facilitation 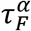 for groups 1-4 was taken from ref.^38^. The time constant of recovery from desensitization, *τ_D_*, was set equal to the value reported in ref.^37^ for all groups, and the parameters *Δ_D_* was chosen so as to obtain the relative reduction in quantal size reported in the same reference^37^. To qualitatively account for the slow approach to steady state transmission observed in MF-GC synapses^37,38^ we set *p_ref_* to a value of 0.6 for all synapse types.

To set the presynaptic quantal parameters we firstly required that the model quantal parameters, *q_0_, N* and *p*, match the average of those measured in ref.^27^ for each synapse group. The estimation of the experimental values 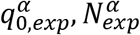 and 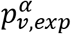 was carried out via multiple-probability fluctuation analysis^27^, which assumes a single vesicle pool. To constrain the corresponding parameters of our two-pool model, we assumed:

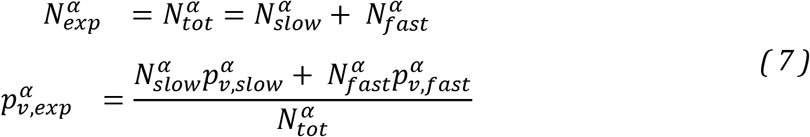

while keeping 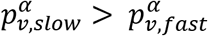. Since the quantal size did not significantly differ between groups^27^, we set *q*_0_ = 1 for all groups for simplicity. As group 4 featured almost no STP, we modeled these synapses without slow pool.

The above equations do not have a unique solution. In order to constrain the synaptic parameters further, we additionally required that the relative unitary response current amplitudes between synapse groups and their pair pulse ratios approximately equal the experimentally measured ones. To account for the fact that group 5’s pair pulse ratio is larger than one, we set *τ_F_* = 30 ms for this group, as in ref.^27^.

Finally, we extracted the relative occurrence of each synapse type from ref.^27^.

A set of synaptic parameters that reproduces the behavior of the 5 synapse groups from ref.^27^ which we used in **Figs. 1, 2** and **6** is summarized in the following table:

|  | Group 1 | Group 2 | Group 3 | Group 4 | Group 5 | Ref. |
| --- | --- | --- | --- | --- | --- | --- |
| $N_{slow}$ | 4 | 3 | 4 | - | 3 | 27 |
| $N_{fast}$ | 16 | 12 | 6 | 10 | 12 | 27 |
| $p_{v,slow}$ | 0.9 | 0.8 | 0.4 | - | 0.4 | 27 |
| $p_{v,fast}$ | 0.72 | 0.55 | 0.35 | 0.3 | 0.15 | 27 |
| $\tau_{ref}^{slow}$ [ms] | 2000 | 2000 | 2000 | - | 2000 | 37 |
| $\tau_{ref}^{fast}$ [ms] | 20 | 20 | 20 | 20 | 20 | 37 |
| $\tau_F$ [ms] | 12 | 12 | - | 12 | 30 | 27,38 |
| $p_{ref}$ | 0.6 | 0.6 | 0.6 | - | 0.6 | - |
| $\Delta_D$ | 0.1 | 0.1 | 0.1 | 0.1 | 0.1 | 37 |
| $\tau_D$ [ms] | 100 | 100 | 100 | 100 | 100 | 37 |
| occurrence | 6% | 16% | 36% | 24% | 16% | 27 |

### MF firing rate parameters

MF firing rate distributions of the full CCM_STP_ were set according to the broad range described in the literature^71,72,65,66,73,40,74–77^. MFs forming synapse types 1 and 2, which convey primary sensory information, were set to high firing frequencies according to experimental observations^40,72^ (see Fig. 1b, left panels), while those firing rates for the other synapses types were lower^75,76^. For the full model, this led to synapses with high *p_v_* being associated with MF inputs with comparatively higher average firing rates (primary sensory groups 1, 2) and synapses with low *p_v_* being associated with MF inputs with comparatively lower average firing rates (secondary/processed sensory groups 3, 4, 5). We chose to describe MF firing rate distributions by Gaussian distributions whose negative tails were set to zero. Means and standard deviations of the Gaussian distributions were set such that the means and standard deviations of the resulting thresholded distributions resulted in the values summarized in the following table:

|  | Group 1 | Group 2 | Group 3 | Group 4 | Group 5 |
| --- | --- | --- | --- | --- | --- |
| $\mu$ [Hz] | 200 | 200 | 20 | 20 | 20 |
| $\sigma$ [Hz] | 20 | 20 | 20 | 20 | 20 |

### Cerebellar cortical circuit model

The standard cerebellar cortex model with STP (CCM_STP_) consists of firing rate units representing *M* = 100 MFs, *N* = 3000 GCs, a single PC, and a single molecular layer interneuron (MLI), i.e. each neuron’s activity is represented by a single continuous value corresponding to an instantaneous firing rate. The PC is spontaneously active (40 Hz) and linearly sums excitatory inputs from GCs and inhibition from the MLI. Each GC received four MF synapses, randomly selected from the different synapse types according to their experimentally characterized frequency of occurrence^27^. The synaptic inputs to the GCs and their firing rates are given by:

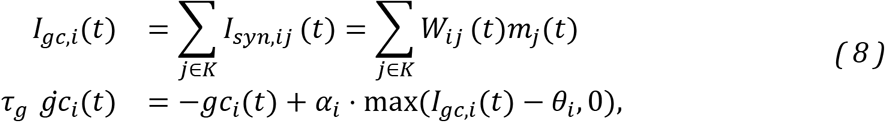

Where the granule cell membrane time constant *τ_g_* = 10ms. In the above equation, *K* is a set of four indices, randomly drawn from the indices of all MF with the constraint that at least one of these stems from a MF belonging to group 1, 2 or 5, as observed experimentally^27^. The gain *α_i_*, and threshold *θ_i_* are set individually for each GC *i* as explained below.

MLI activity is assumed to represent the average rate of the GC population, thus allowing each GC to have a net excitatory or inhibitory effect depending on the difference between the MLI-PC inhibitory weight and the respective GC-PC excitatory weight:

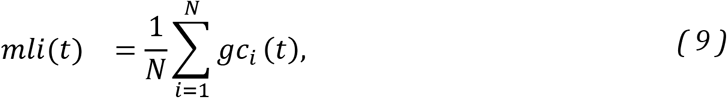

The synaptic weights between the *i*th GC and the PC and between the MLI and PC were defined as *J_E,i_* and *J_I_*, respectively. The total synaptic input to the PC is thus given by:

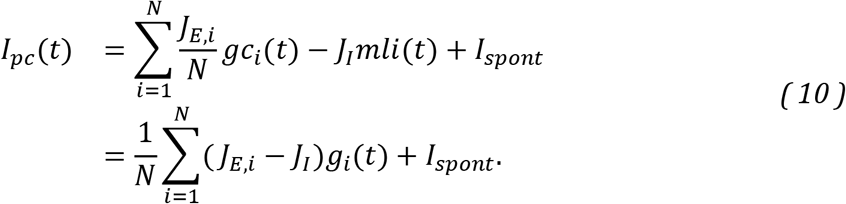

Here, *I_spont_* is an input that maintains the spontaneous firing of the PC at 40 Hz.

Finally, the PC firing rate is given by:

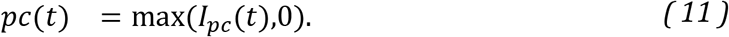

In **Fig. 1**, the GC-PC weights *J_E,i_* were drawn from an exponential distribution with mean equal to 1. In order to decrease or increase the ratio of the average excitatory to inhibitory weight in **Fig. 1c** and **1d**, we set *J_I_* = 1.025 and *J_I_* = 0.975 respectively. The synaptic, full CC model, and the reduced model described below were numerically integrated using the Euler method with step size 0.5 ms.

#### GC Threshold and gain adjustment

Changing the statistics of the MF firing rate distributions changes the fraction of active GCs at any given time and the average GC firing rates. To avoiding the confounding impact that co-varying these quantities has on learning performance when comparing different MF parameter sets we adjusted GC thresholds, *θ_i_* and gains *α_i_* such that, at steady state, the fraction of active GCs and the average GC firing rates were identical for all MF parameter choices. Specifically, we drew 1000 random MF patterns from the respective firing rate distributions and we calculated the steady inputs values of the synaptic dynamics as follows:

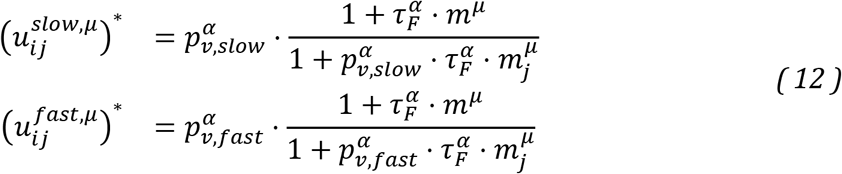

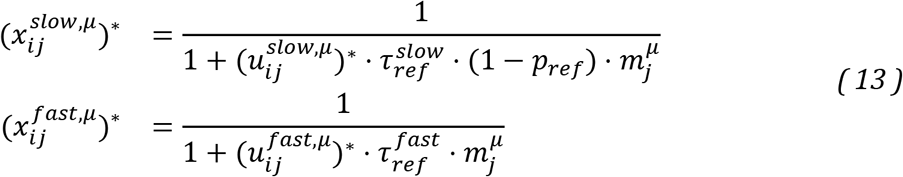

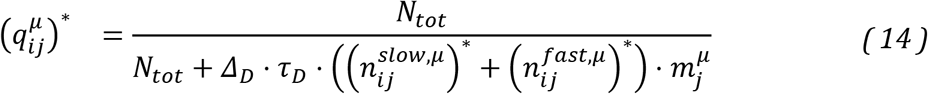

With these, we obtained, for each individual GC, the distribution of steady state inputs and firing rates:

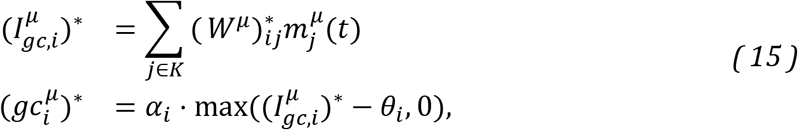

We then adjusted *α_i_* and *θ_i_* such that, for each GC, the average steady state GC rate was 5Hz across all patterns and the fraction of patterns with non-zero GC activity was 0.2 which is within the range of experimental observation^77,78^. Throughout the article, this adjustment was carried out every time we changed synaptic parameters (Fig. 5), the parameters of the MF firing rate distributions (Fig. 4) or the MF to synapse connectivity (Fig. 5).

#### Supervised learning rule

Eyelid conditioning learning was implemented by adjusting the *J_E,i_* via a supervised learning rule. For a given delay *t_target_* we used the following target signal to produce PC pauses:

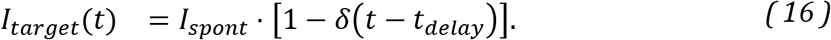

where *δ*(*x*) = 1 if *x* = 0 and *δ*(*x*) = 0 otherwise. The target PC firing rate *I_target_* (*t*) is a Dirac pulse in which the PC rate is zero at a particular time *t_target_* following the start of the CS. We quantify the deviation of the PC firing rate from the target rate by the least-squares loss *E* which is to be minimized during learning:

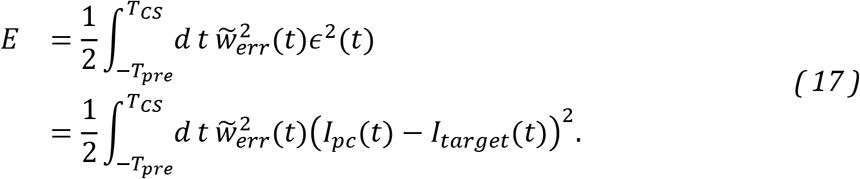

Here, *T_CS_* is the time interval after CS onset during which we require the PC to follow the target signal and *T_pre_* a time interval before CS onset during which the PC should fire at its spontaneous rate. *ϵ*(*t*) denotes the deviation between target and actual PC output at time *t*. 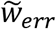 is a weight factor that we use to increase the impact of the target time, given by:

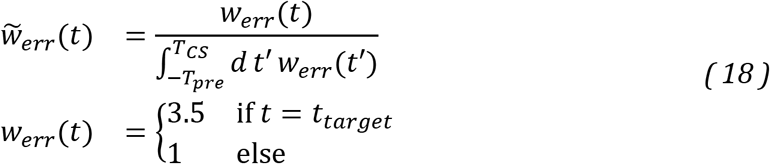

In all main figures we used *T_CS_* = 1.4*s* and *T_pre_* = 0.1*s*.

GC-PC weights *J_E,i_* were modified during learning using gradient descent to reduce the error *E* at each step of the learning algorithm:

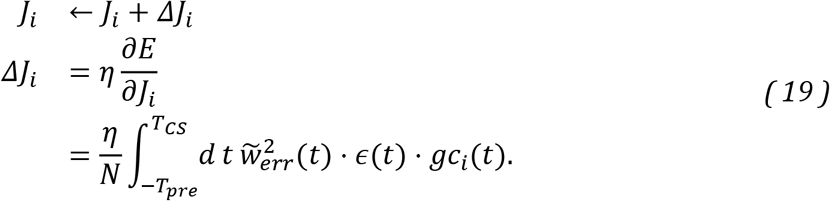

Here, *η* is a learning rate. For our simulations, we modified this basic rule in two ways. Firstly, similar to ref.^79^, we explicitly simulated a climbing fiber (CF) rate, *cf*, that is modulated by the error signal *ϵ*(*t*) = *I_pc_* (*t*) – *I_target_* (*t*) according to:

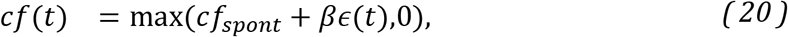

where *cf_spont_* is the spontaneous CF rate and *β* a proportionality factor. The CF rate was then used to obtain the following modified the weight update:

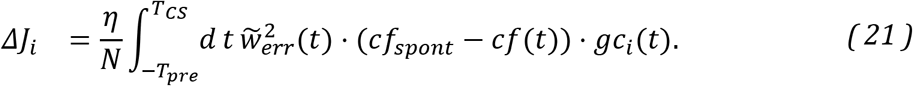

where we also set *J_E,i_* = 0 when a learning iteration resulted in a negative weight. As the CF rate is required to be positive or zero, this formulation limits the error information that is transmitted to the PC as compared to the simple gradient rule. Note that this learning rule yields synaptic long-term depression when CF and GC are simultaneously active and long-term potentiation when GC are active alone, consistent with experimental data on GC-PC synaptic plasticity^55^.

Furthermore, recent experimental findings suggest that temporal properties of GC-PC plasticity rules are tuned such as to compensate for the typical delays expected for the time for error information to arrive in the cerebellar cortex^69^. Here, we did not explicitly model CF error information delays, and for the sake of simplicity, directly modeled the timing of PC activity to show that the GC basis set is sufficient to generate an appropriately timed PC pause.

To increase the learning speed, we added a Nesterov acceleration scheme to equation (21)^80^, introducing a momentum term to the gradient, making weight updates during a given iteration of the algorithm dependent on the previous iteration. The implementation we chose additionally features an adaptive reset of the momentum term, improving convergence properties^80^. This addition does not reflect biological mechanisms and is used mainly for practical convenience.

We used simulated GC activity that was subsampled by a factor of 10 and set *η* = 0.0025, *β* = 0.5 and the initial distribution of weights to *J_E,i_* = *J_I_* = 10 for all *i*. For all eyelid response learning simulations, we chose *cf_spont_* = 1 *Hz* (**Figs. 2, 4, 5**).

#### Error measure of learned Purkinje cell pause

We defined the error between the PC pause and the *I_target_* (see Figs 4, S3, S4 and S5) in the following way:

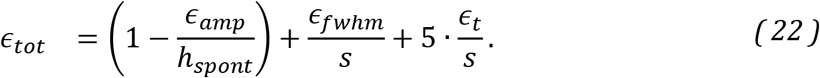

The first term depends on the amplitude of the PC pause relative to baseline firing, yielding a small error when the amplitude goes to zero. The second term corresponds to the normalized width of the PC pause. Finally, the third term is the normalized deviation of the pause’s minimum from the target time, *ϵ_t_*. To increase the importance of this term we scaled it by a factor 5. The error measure in Figs. S4 and S5 is the sum of *ϵ_tot_* over all tested delays.

### Reduced CC model

In the reduced synaptic model, we neglect facilitation and desensitization yielding constant release probabilities and normalized quantal size:

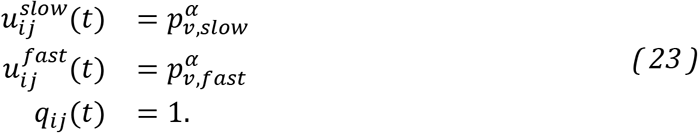

We obtain for the vesicle pool dynamics:

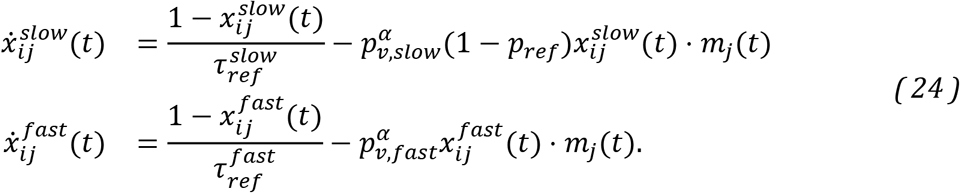

and the total synaptic weight becomes:

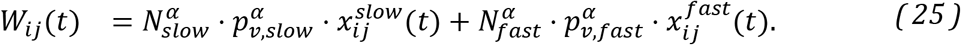

Here the index *α* denotes membership in the driver or supporter category. The synaptic currents of the reduced model are computed as in the full model. Each GC receives exactly two driver and two supporter MF inputs with random and pairwise distinct identities. To eliminate any non-synaptic dynamics from the reduced model, we removed the GC membrane time constant yielding GC dynamics that follow the synaptic input instantaneously:

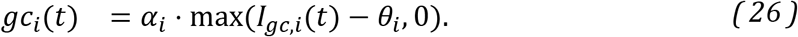

Finally, GC threshold and gain adjustments were carried out as for the full CCM_STP_ where instead of equations (12) and (14) we used equations (23).

#### Synaptic parameters of the reduced model

The parameters of the reduced model we set to create two synapse types that capture the essence of the experimentally observed synaptic behavior: a strong and fast driver synapse, and a weak and slow supporter synapse. All synaptic parameters of the model used in **Fig. 3**, **4** and **6** are summarized in the following table:

|  | Drivers | Supporters |
| --- | --- | --- |
| $N_{slow}$ | 3.5 | 4 |
| $N_{fast}$ | 14 | 6 |
| $p_{v,slow}$ | 0.8 | 0.4 |
| $p_{v,fast}$ | 0.6 | 0.2 |
| $\tau_{ref}^{slow}$ [ms] | 2000 | 2000 |
| $\tau_{ref}^{fast}$ [ms] | 20 | 20 |
| $p_{ref}$ | 0.6 | 0.6 |
| occurrence | 50% | 50% |

In **Fig. 5**, firing rates and release probabilities were randomly drawn from uniform distributions. In detail, the release probabilities of the slow pool, *p_v,slow_*, were drawn from distributions with a lower and upper bound of 0.1 and 0.9, respectively, (**Fig. 5a, d, g** and **h**), and the corresponding release probabilities of the fast pool were calculated according to 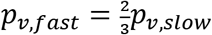, keeping them strictly lower. Lower and upper bounds of the distribution of firing rates used in panels **a** and **d** were 5 Hz and 270 Hz, resulting in firing rate standard deviations of *σ_rate_* ≈ 38.2 Hz for the two-groups case (**Fig. 5a**) and *σ_rate_* ≈ 15.3 Hz for the five-groups case (**Fig. 5d**). The bounds of the distributions in panels **g** and **h** were chosen such as to produce average group firing rates equal to those in panel **d** and firing rate standard deviations that increased with the group index, i.e. *σ_rate_* ≈ {5.0, 7.6,10.2,12.7,15.3} Hz for groups 1 to 5, respectively. Finally, the sizes of the slow vesicle pool were fixed at *N_slow_* = 4 and the size of the fast vesicle pools were set to decrease with the group index, i.e. *N_fast_* = {16,6} for the two-groups case, and *N_fast_* = {16,12,8,6,6} for the five-groups case. Finally, the desired rank correlation between *p_v_* identities and MF identities was achieved by creating a Gaussian copula reflecting their statistical dependency and reordering the marginal *p_v_* and MF distributions accordingly.

### Derivation of *τ_syn_* and *A_t_*

In the reduced model, we derived an analytical solution to the synaptic current driving a GC in response to the CS. Since the equations describing slow and fast vesicle pool dynamics are formally very similar, we describe the derivation for a single slow pool only. Additionally, we suppress all indices for the sake of readability. We assume that the MF rate *m(*t**) switches instantaneously from *v_preCS_* to *v_CS_* at time *t*′ = 0. Integration of equations (24) from *t*′ = 0 to *t* yields:

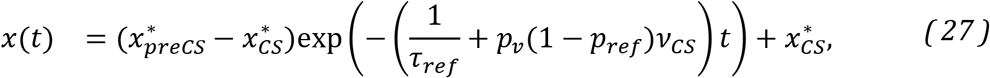

Here, 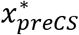 and 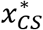 denote the steady state values of x before (preCS) and after (CS) the firing rate switch. They are given by:

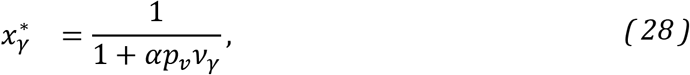

with

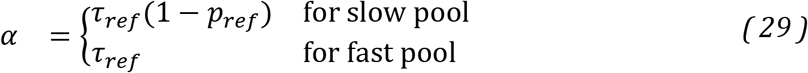

From equation (27) we can read off the synaptic time constant that governs the speed of transition from steady state a value before the CS to a steady state during the CS:

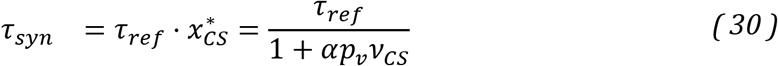

This equation is similar to one derived previously^51,81^. The total synaptic current per unit time for a single pool during the CS is given by:

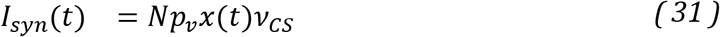

Combining equations (27) and (31) we get:

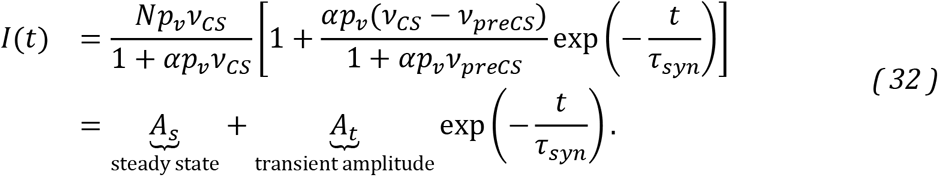

Thus, the transient amplitude for a single vesicle pool is:

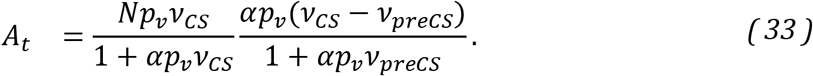

For a single synapse, the total transient amplitude is the sum of the individual fast pool and slow pool transients:

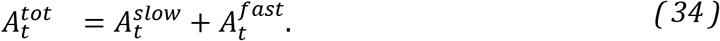

To generate the surface plots in **Fig. 4** and **Fig S3** we generated 10^5^ firing rates from the driver and supporter MF rate distributions, respectively, and used equations (30), (33) and (34) to calculate the corresponding values of the *A_t_* and *τ_syn_*. From these, the plots of the joint *A_t_* and *τ_syn_* distribution and the marginal distributions were generated using a two- or one-dimensional kernel density estimator, respectively^82^. Note that, formally, *τ_syn_* is maximal when *v_CS_* = 0. In that case, however, there is no synaptic transmission as 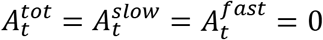. When plotting the joint *A_t_-τ_syn_* distribution in **Fig. 4** and **Fig S3**, we therefore omitted time-constants and transient amplitudes corresponding to *v_CS_* = 0.

### Bayesian estimation of time intervals

In CCM_STP_ simulations, the “ready” signal was treated as an instantaneous switch of the MF input rates that persisted over the course of a trial (**Fig. 6c**), similar to the CS in the eyelid learning simulations. To learn a distribution of ready-set intervals, we generated target signals within intervals that varied from iteration to iteration of our learning algorithm (**Fig. 6c**). We performed simulations with the following five different uniform distributions of ready-set intervals: 25-150 ms, 50-200 ms, 100-300 ms, 200-400 ms, 300-500 ms. Learning was carried out separately for each interval and for 12000 iterations. We found that in order to achieve the correct biases for the two longest intervals, we had to introduce a higher CF baseline firing rate, *cf_spont_* = 5 Hz, possibly indicating a higher importance of LTP for this task. The other learning parameters were kept the same as in the eyelid learning simulations.

In keeping with ref.^30^, we modeled the DN neuron an integrator, whose rate was calculated according to:

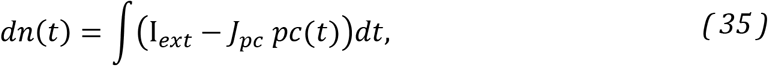

where the *J_pc_* is the weight of the inhibitory PC-DN synapse and I*_ext_* = 〈*pc*〉 is an external excitatory input to DN. It was set equal to the average PC firing rate during the ready-set period in order to ensure that excitation and inhibition onto the DN are of comparable size. For simplicity we set *J_pc_* = 1.

In order to map the DN rate to a time axis (**Fig. 6f,j**), we rescaled every individual DN output curve according to:

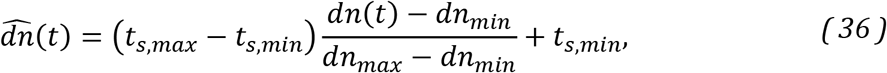

where *t_s,max_* and *t_s,min_* are the maximum and minimum of the respective ready-set interval and *dn_max_* and *dn_min_* are the maximum and minimum values of the DN firing rate. Since the transformation (36) is linear, the essential features exhibited by the DN firing rate (i.e. its biases) are preserved.

To show how the theoretical Bayesian least squares (BLS) interval estimate can be obtained, we follow the reasoning from ref.^30^. It is assumed that, in the ready-set-go task, subjects perform a noisy measurement, *t_m_*, of the ready-set interval, *t_s_*, according to:

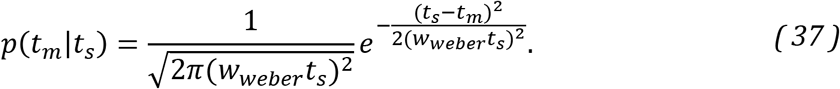

Note that the standard deviation of the estimate of *t_m_* increases with the length of the interval *t_s_* with proportionality factor *w_weber_*, which is the weber fraction. Given the distribution of time intervals in the ready-set-go task, Π(*t_s_*), the Bayesian estimate of *t_s_* given *t_m_* is:

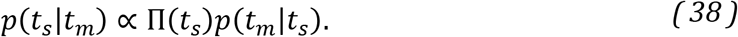

The BLS estimate is the expected value of the previous expression:

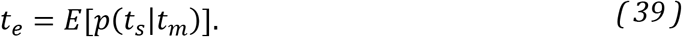

We performed a least-squares fit of the BLS model to the CCM_STP_ outputs (from all five interval distributions simultaneously) with *w_weber_* as a single free parameter.

### Recurrent Golgi cell inhibition

To probe the effect of recurrent inhibition in the reduced CCM_STP_, we added one Golgi cell (GoC) that received excitatory inputs from all GCs and formed inhibitory synapses onto all GCs. For simplicity, we assumed that the GoC fires with a rate *goc* equal to the average GC firing rate, similarly to the MLI, and that all GoC to GC synapses have identical weights, *J_goc_*:

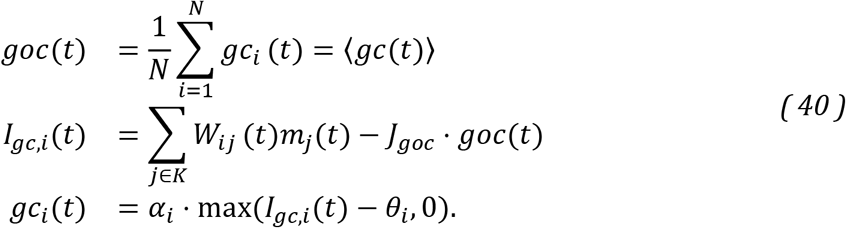

The above equations imply that, in this configuration, the GoC acts as an activity-dependent GC threshold.

To ensure that the overall GC activity level in the reduced CCM_STP_ with GoC inhibition is comparable to the case without, we require the same criterion as above: an average GC rate of 5Hz and a fraction of activated GCs of 0.2 in steady state. Since the average GC input now depends on the average GC firing rate itself, manual adjustment of GC thresholds, *θ_i_*, and gains, *α_i_*, carried out above, is not feasible.

Instead, a steady-state solution of equations (40) satisfying our requirements has to be found numerically. We first set up the CC network without the GoC and adjusted GC thresholds, *θ_i_*, and gains, *α_i_*, according to the procedure described above. Note that in the reduced model, due to every GC receiving the same combination of inputs (i.e. 2 supporter and two driver inputs), both *θ_i_* and *α_i_* are similar across GCs. We thus made the additional simplification of setting *θ* = E(*θ_i_*) and *α* = E(*α_i_*) for all GCs. We then reduced GC thresholds by 10% and introduced the GoC.

To obtain the average steady state GC firing-rate we assumed that the synaptic currents of a single GC are normally distributed across MF input patterns, or, equivalently, across GCs. Mean and variance of the GC inputs are:

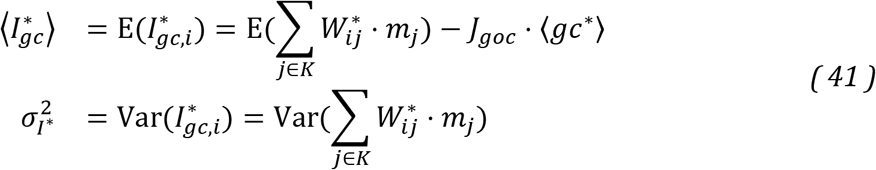

We can then express the average GC firing rate in the *N* → ∞ limit as:

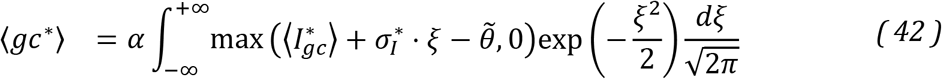

where 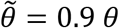. The fraction of active GCs *f* can be written as:

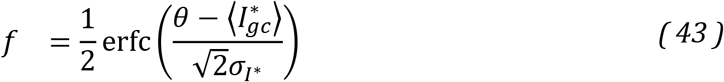

We can now impose that

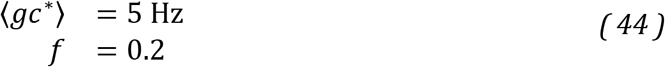

and find a self-consistent solution of equations (41), (42) and (43) by adjusting the parameters *J_goc_* and *α*. To do so we used the hybrid numerical root-finder from the GNU scientific library^83^ with default step size.

### Software implementation

Figures were generated using Matlab. All simulations were performed using custom C++ code and are available on a GitHub repository ().

## Supplementary Figures

**Supplementary Figure 1.**
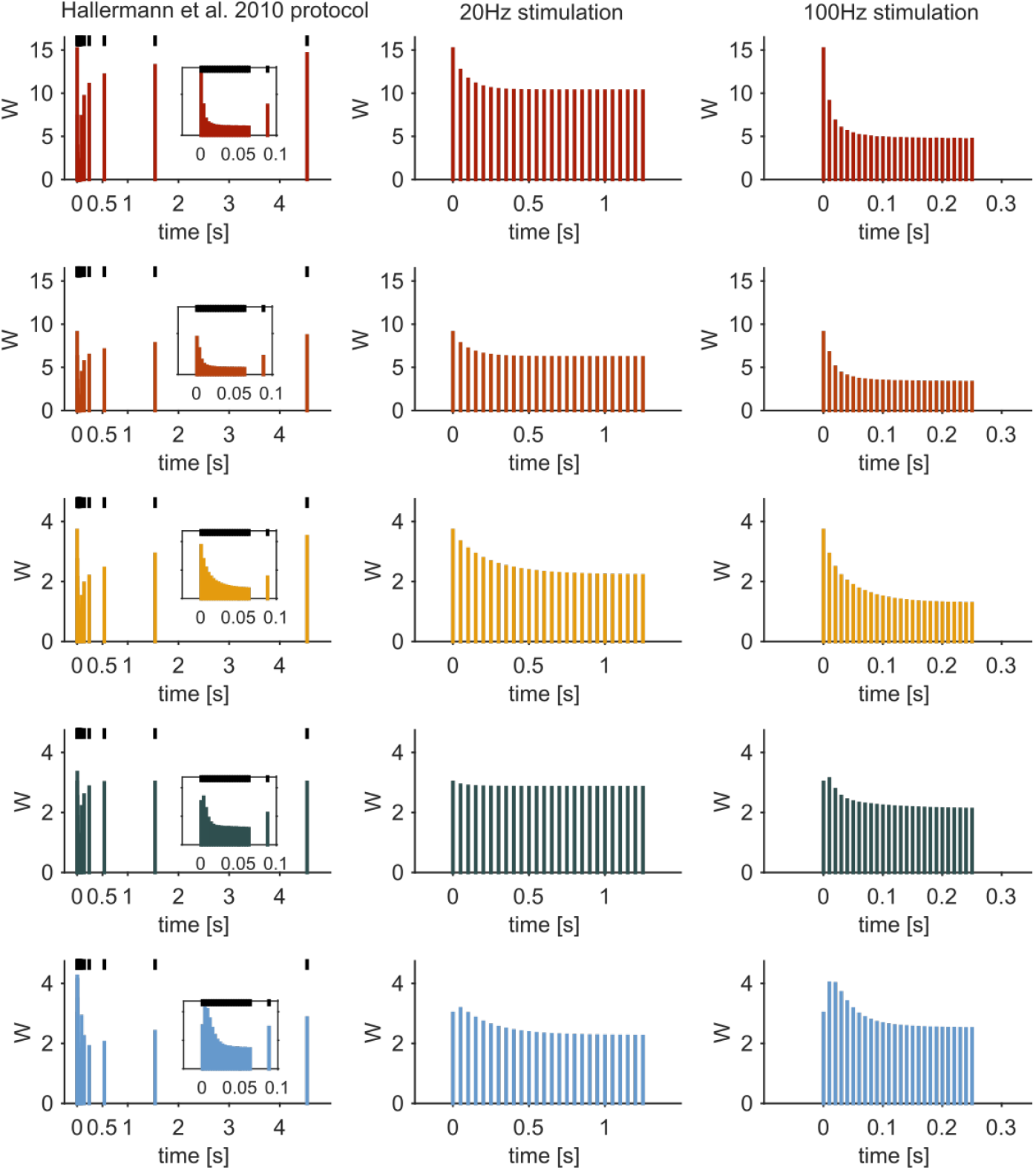
Behavior of model synapses subjected to different stimulation protocols. Each row represents one of the five synapse types from **Fig. 1**. First column: average synaptic weight in response to 300 Hz train followed by increasing intervals ranging from 25 ms to 5 s as in ref.^37^. Inset: zoom on 300 Hz train. Second and third column: average synaptic weight in response to trains of 26 stimuli at 20 Hz and 100 Hz, respectively, similar to ref.^38^

**Supplementary Figure 2.**
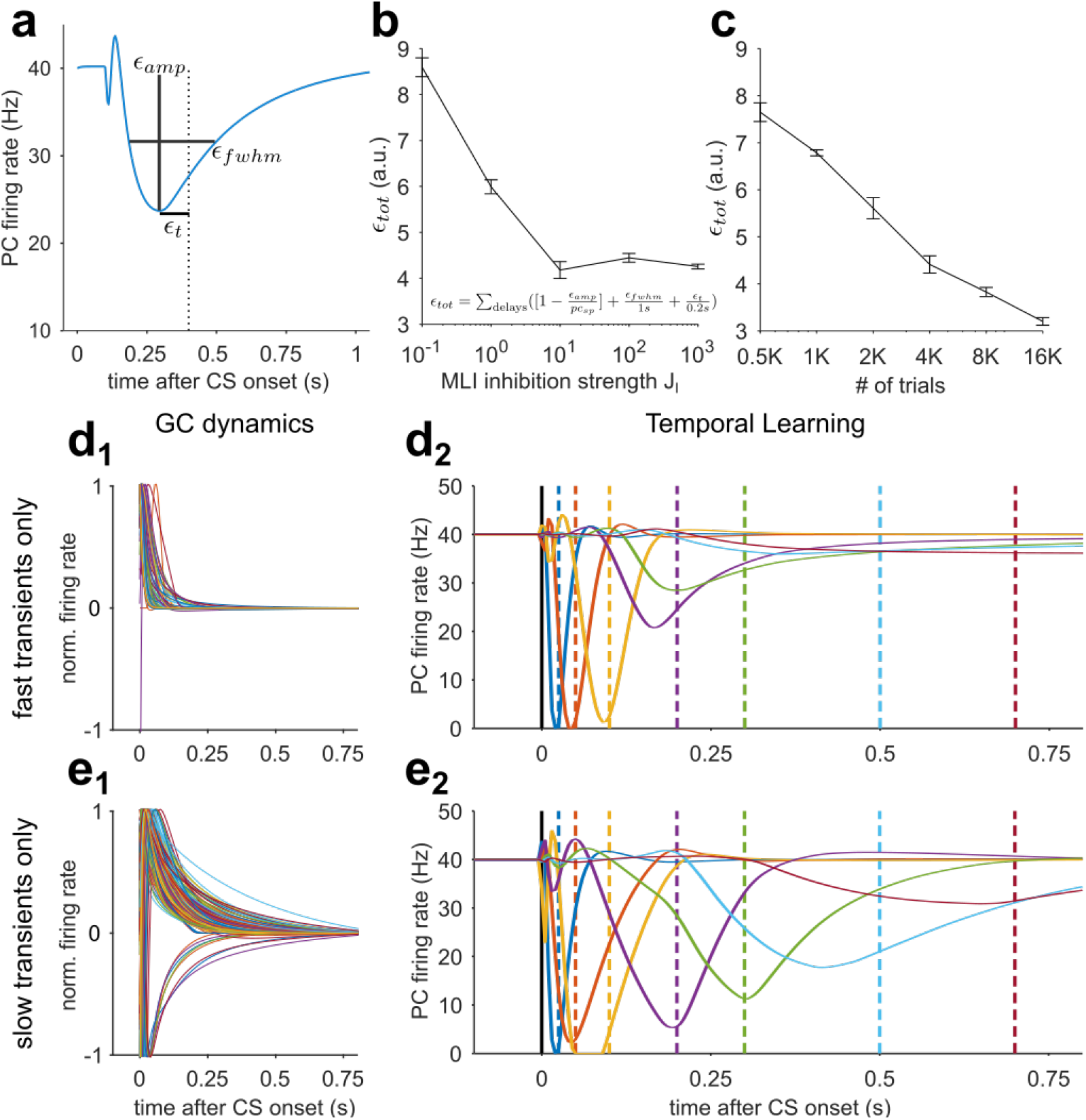
***(a)** Definition of error components used to assess learning performance. **(b)** Learning error as a function of MLI-PC inhibition strength. The total error is computed from the individual components according to the formula shown. Black line represents the average over 20 realizations of the network in **Fig.2**. Error bars are SEM. **(c)** Same as **b** for learning error as a function of number of learning iterations. **(d_1_)** As in **Fig.2e**, but without GCs whose transients decayed to 10% of their peak values in more than 150 ms. **(d_2_)** Learning performance when using the temporal basis from panel **d_1_**. **(e_1_)** As in **Fig.2e**, but without GCs whose transients decayed to 10% of their peak values in less than 150 ms. **(e_2_)** Learning performance when using the temporal basis from panel **e_1_***. Note that learning of short delays was not completely abolished because our manipulations did not affect the rising phases of the GC transients.

**Supplementary Figure 3.**
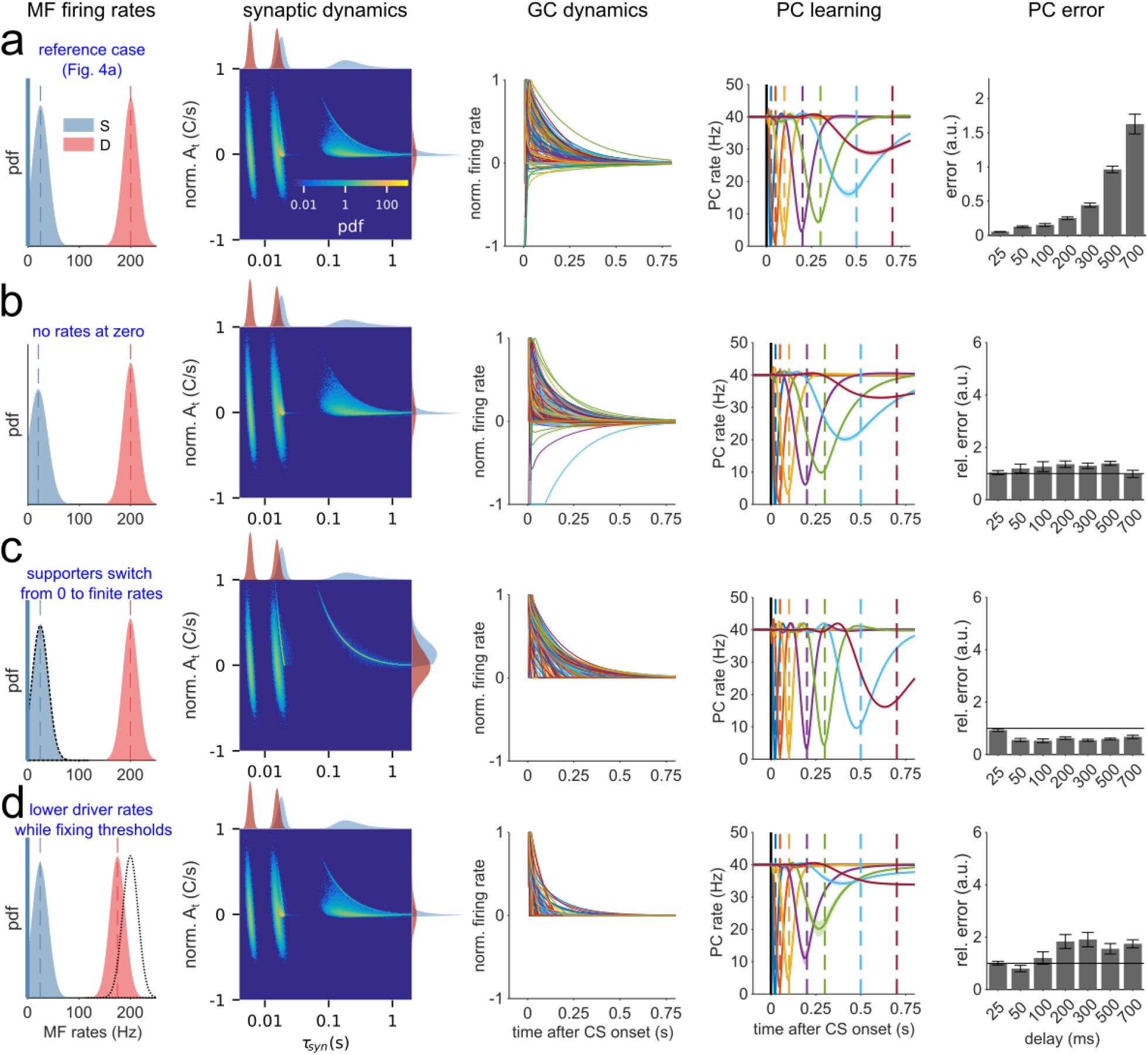
Additional examples of MF rate distributions and their impact on learning **(a)** Same as **Fig. 4a** (i.e. reference case). **(b)** Without zero firing rates. **(c)** Simulations in which supporter inputs are set to zero before the CS and switch to finite firing rates during the CS. **(d)** Simulations in which GC thresholds are set as in **a** (blue and dashed firing rate distributions), but eyelid conditioning is carried out with lower driver inputs (blue and red firing rate distributions).

**Supplementary Figure 4.**
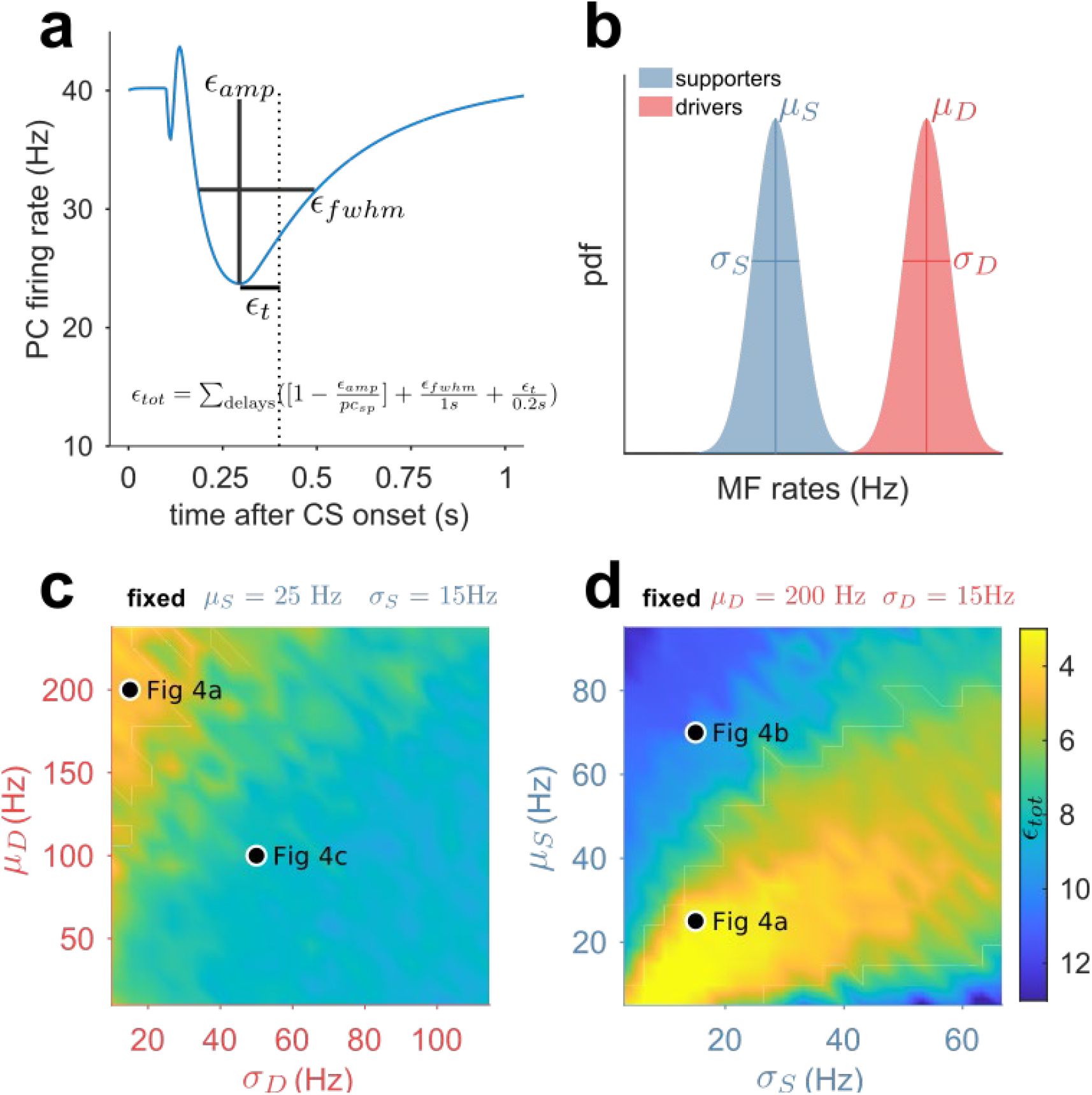
Scan of MF firing rate parameters show distinct roles for driver and supporter inputs **(a)** Definition of error components used to assess learning performance. The total error is computed from the individual components according to the formula shown. **(b)** Definition of the MF firing rate parameters. μ_D_, μ_s_ and σ_D_, σ_s_ denote means and standard deviations of the MF firing rate distributions of drivers (red) and supporters (blue) respectively. **(c)** Scan over driver parameters while keeping supporter parameters fixed. **(d)** Scan over supporter parameters while keeping driver parameters fixed. In **c** and **d** the total learning error is color coded and parameter configurations corresponding to rows in **Fig. 4** are indicated by black dots.

**Supplementary Figure 5.**
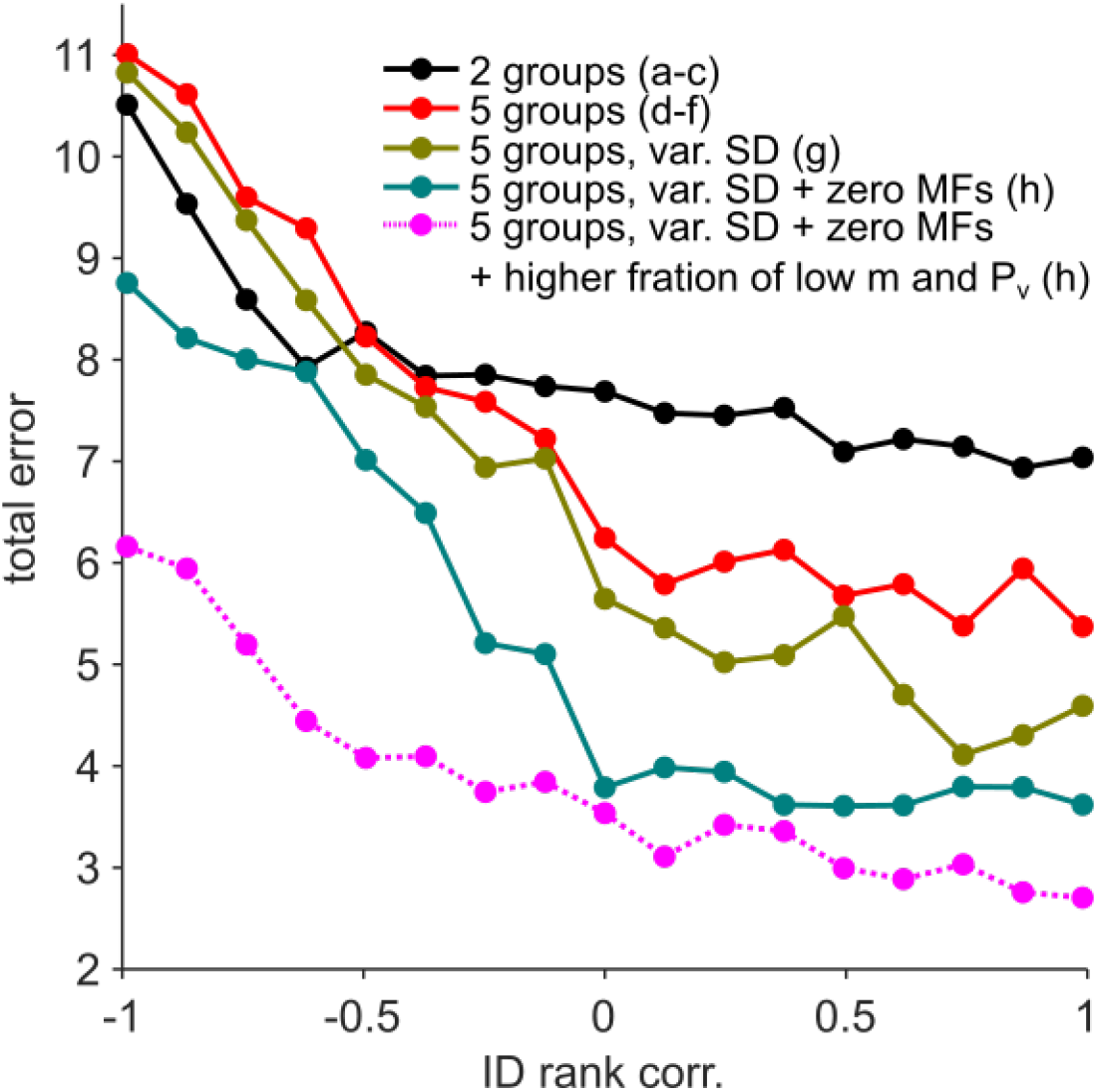
Average total error over seven delay intervals for varying rank-correlation between the m category and the p_v_ category for all scenarios shown in **Fig. 5**.

**Supplementary Figure 6.**
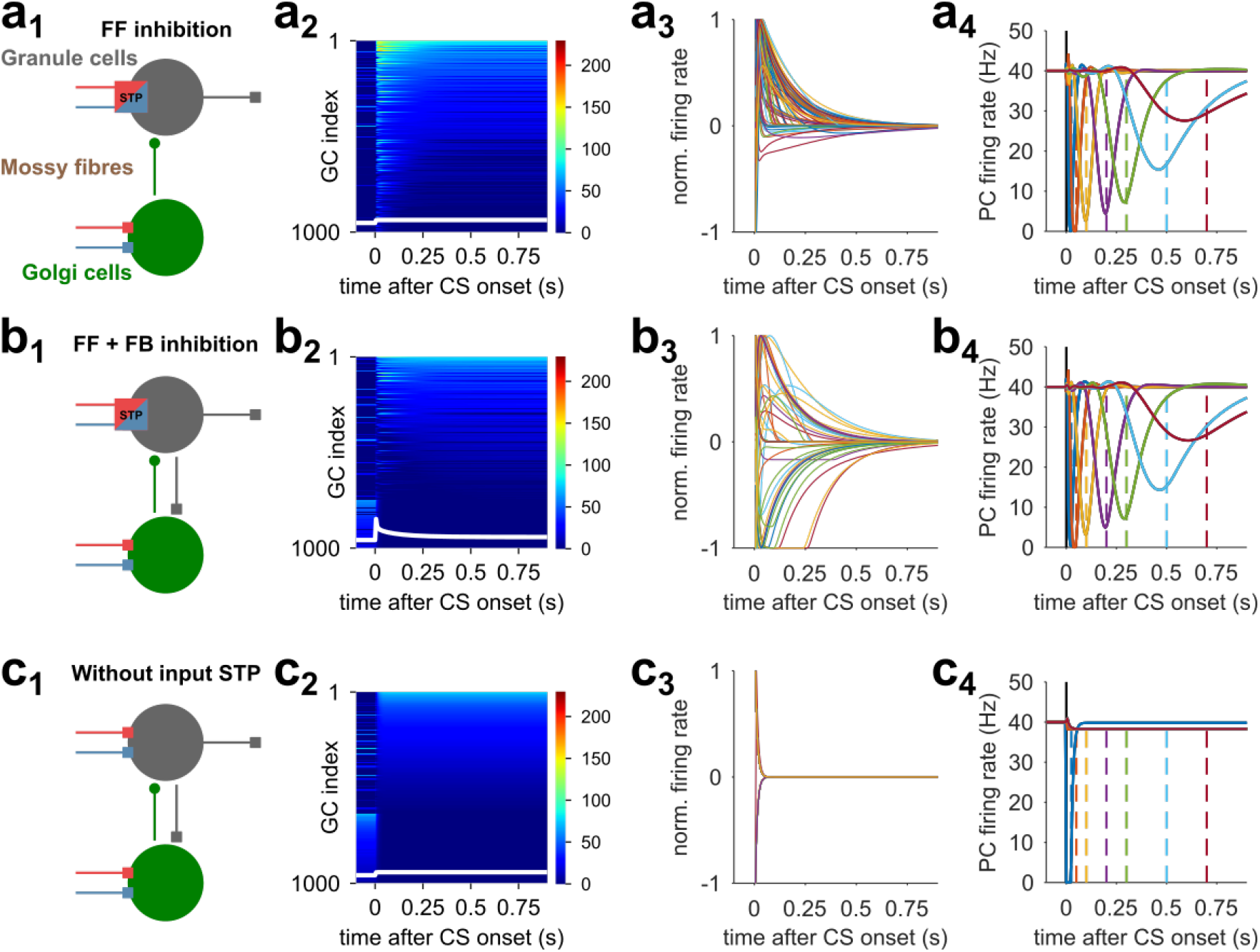
Eyelid learning is not significantly affected by simple GoC feedback **(a_1_)** Scheme of CC input layer with GoC inhibition that acts in a purely feed-forward manner. This configuration is functionally identical to the reduced model used in the main text. **(a_2_, a_3_)** GC temporal basis (as in **Fig. 2c**). White line in a_2_ indicates GoC activity. **(a_4_)** PC eyelid response learning. **(b_1-4_)** Same as row **a** but with GC-GoC feedback connections. **(c_1-4_)** Same as row **b** but without MF-GC STP.

**Supplementary Figure 7.**
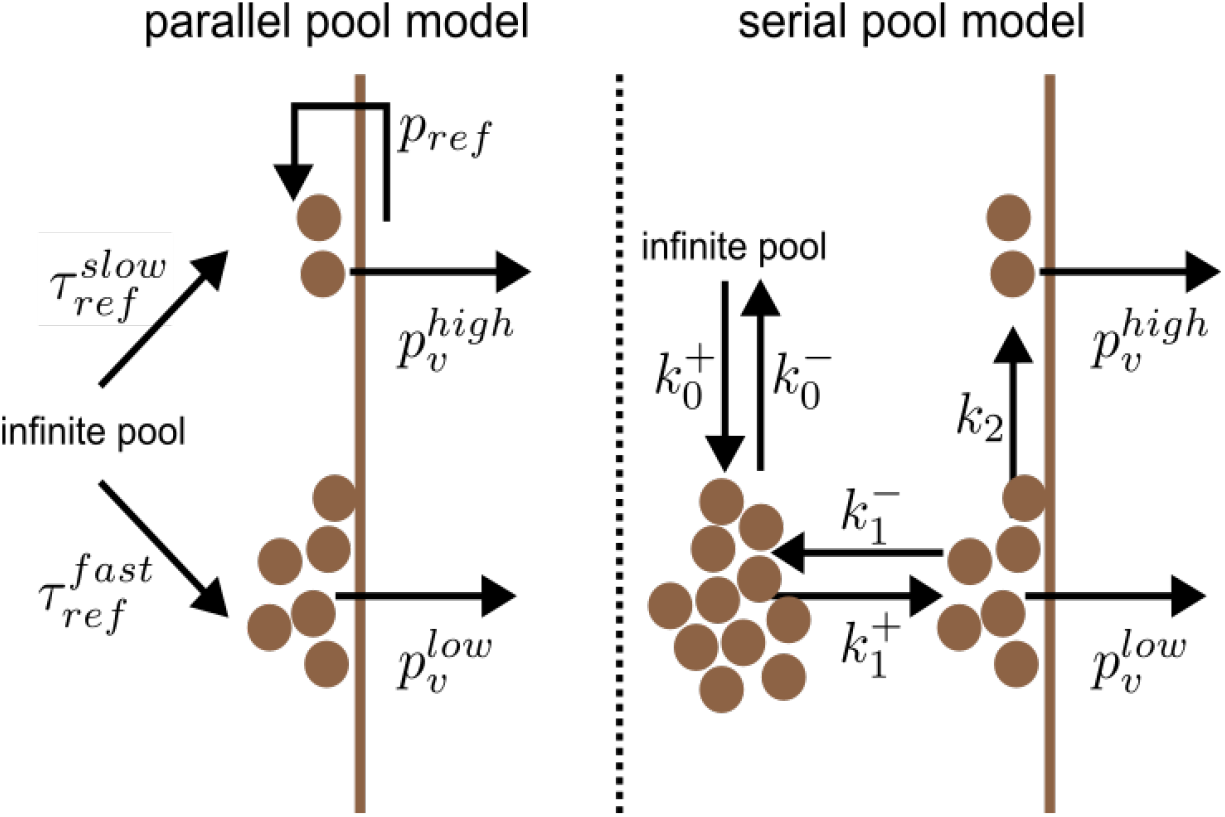
Scheme comparing the parallel pool model used here with the serial pool model from ref.^37^. The symbols 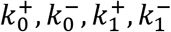 and k_2_ are rate constants described in ref.^37^.

## Notes

### Competing Interest Statement

The authors have declared no competing interest.

